# Robust Decoding of Speech Acoustics from EEG: Going Beyond the Amplitude Envelope

**DOI:** 10.1101/2025.10.07.680996

**Authors:** Alexis D MacIntyre, Clément Gaultier, Tobias Goehring

**Affiliations:** MRC Cognition and Brain Sciences Unit, University of Cambridge, 15 Chaucer Rd, Cambridge CB2 7EF; Université Paris Cité, Institut Pasteur, AP-HP, INSERM, CNRS, Fondation Pour l’Audition, Institut de l’Audition, IHU reConnect, F-75012 Paris, France

**Keywords:** Neural Decoding, Speech Perception, EEG, spectral features

## Abstract

**Objective:** During speech perception, properties of the acoustic stimulus can be reconstructed from the listener’s brain using methods such as electroencephalography (EEG). Most studies employ the amplitude envelope as a target for decoding; however, speech acoustics can be characterised on multiple dimensions, including as spectral descriptors. The current study assesses how robustly an extended acoustic feature set can be decoded from EEG under varying levels of intelligibility and acoustic clarity.

**Approach:** Analysis was conducted using EEG from 38 young adults who heard intelligible and non-intelligible speech that was either unprocessed or spectrally degraded using vocoding. We extracted a set of acoustic features which, alongside the envelope, characterised instantaneous properties of the speech spectrum (e.g., spectral slope) or spectral change over time (e.g., spectral flux). We establish the robustness of feature decoding by employing multiple model architectures and, in the case of linear decoders, by standardising decoding accuracy (Pearson’s *r*) using randomly permuted surrogate data.

**Main results:** Linear models yielded the highest *r* relative to non-linear models. However, the separate decoder architectures produced a similar pattern of results across features and experimental conditions. After converting *r* values to Z-scores scaled by random data, we observed substantive differences in the noise floor between features. Decoding accuracy significantly varies by spectral degradation and speech intelligibility for some features, but such differences are reduced in the most robustly decoded features. This suggests acoustic feature reconstruction is primarily driven by generalised auditory processing.

**Significance:** Our results demonstrate that linear decoders perform comparably to non-linear decoders in capturing the EEG response to speech acoustic properties beyond the amplitude envelope, with the reconstructive accuracy of some features also associated with understanding and spectral clarity. This sheds light on how sound properties are differentially represented by the brain and shows potential for clinical applications moving forward.

## 1. Introduction

Neural decoding is a form of analysis whereby properties of a presented stimulus are reconstructed from an observer’s neural activity. When framed as a system identification problem, the objective is to estimate an output signal (the perceived stimulus) given the state of the input (measured brain response) [1]. Linear regression-based approaches have proven especially popular thanks to their relative simplicity and reliable performance–even for data with low signal-to-noise ratios (SNRs) obtained using noninvasive methods like electroencephalography (EEG) [2, 3, 4]. Nonetheless, techniques such as mutual information analysis or deep learning-based frameworks are also proposed to capture additional, non-linear components of the stimulus-brain relationship [5, 6, 7]. Although neural decoding is agnostic to any one behavioural domain or modality, the technique has found particular traction within auditory speech perception research, facilitating rapid experimental advancements in topics ranging from auditory attention [8, 9] to infant language acquisition [10]. Yet, a key question is whether neural decoding will prove useful within clinical settings, for instance, to assess hearing device function or pinpoint the source of audiological problems [4, 11]. This promise may rest on aspects of speech decoding that remain unclear: For example, some studies report a positive association between speech decoding accuracy and behaviourally measured comprehension [12, 13, 14, 15]. Other work, however, shows that this relationship is non-contingent on understanding *per se*. Rather, neural differences may arise due to acoustic confounds attributable to the signal processing applied in order to manipulate intelligibility, such as the addition of background noise [16, 17, 18, 19]. One limitation in prior research is that speech decoding studies usually operationalise the acoustic stimulus as its amplitude envelope. The speech envelope is a feature that conveys how signal intensity–roughly equivalent to the percept of loudness–varies over time, and is theorized to be highly salient for understanding [20, 21]. However, the envelope can be reconstructed for unattended speech [5, 22], as well as speech so acoustically altered that phonemic identity is virtually absent [23]. Moreover, recent evidence from older adults with and without hearing loss indicates that envelope decoding may actually be enhanced in these individuals, relative to young and/or typically hearing adults [24, 25]. In sum, envelope decoding may not reliably indicate comprehension, nor optimal, speech-specific processing more generally [26]. The question, thus, arises if there are other acoustic dimensions more suitable to this purpose.

Spectral features, in particular, could provide a more nuanced measure of speech signal conveyance to the brain. Like the amplitude envelope, these features are used to describe time-varying properties of continuous speech, but within the frequency domain. For instance, spectral flux measures how the distribution of energy across frequency bins changes over time, and the spectral centroid provides an estimate of the spectrum’s “centre of gravity”, or its weighted average across frequencies [27]. These and other spectral features may correspond to perceptually relevant speech cues, and thus, serve as useful clinical targets for speech decoding.*‡* To give an example, cochlear implants (CI) work by converting an acoustic input signal into patterns of discrete electrical pulses, which are then used to directly stimulate the auditory nerve via a surgically-implanted electrode array [30, 31, 32]. As most CI contain 12 *−* 24 electrodes, the transmitted signal is considered to have lower spectral resolution than biological hearing, which entails thousands of sensory hair cell receptors. Signal integrity can be further impacted by factors such as cross-channel interference at the electrode-neural interface [33], or non-functioning regions of the auditory nerve [34]. Such complications may further impede sensory access, leading to poorer speech perception outcomes. At present, there are few objective measures available to characterise the auditory signal quality as it reaches the listener’s brain, and individual differences in speech comprehension remain largely unexplained [35]. Hence, speech decoding for spectral features could serve as a candidate biomarker of speech information transmission in CI listeners, especially if found to be sensitive to differences in the spectral resolution of the perceived signal. Incidentally, recent work using invasive electrophysiology in epilepsy patients found that the neural response to spectral flux is sensitive to speech rate and may also reflect phonemic processing; however, that study employed clear and intelligible speech only [36]. Thus, it is unknown how well spectral flux or other spectral descriptors can be decoded under varying levels of, for example, spectral clarity. Moreover, the robustness of spectral feature decoding is currently unestablished. We define robustness as the stability of a feature across different decoding techniques, in that we should expect to find a common pattern of results (e.g., accuracy over participants or listening contexts) regardless of the method employed. Another property of robust decoding is resilience to spurious correlations between the reconstructed feature and its ground truth, given that the corresponding noise floor cannot be known *a priori*. Indeed, in a previous study, we observed that the threshold for decoder significance, which we determined using randomly permuted surrogate data, varied across experimental conditions [19]. If the likelihood of spurious decoding differs across features, the direct comparison of unscaled accuracy scores, typically defined as Pearson’s *r*, would be inappropriate. As an alternative, observed *r* can be compared to a distribution of *r* generated using the same process but from randomised data, thereby providing a measure of decoder confidence.

### The current study

This work investigates the neural decoding of a set of speech acoustic features, including both envelope-based and spectral descriptors, from EEG during a naturalistic listening task. Firstly, we compare acoustic features in terms of the magnitude and robustness of decoding overall, and secondly, we assess which features are most sensitive to two factors relevant for clinical applications: Speech comprehension, which we manipulate linguistically, and acoustic clarity, which we manipulate via a form of spectral degradation called vocoding. We address the robustness of feature decoding by implementing three distinct decoding architectures, including linear and non-linear models; and by accounting for the uncertainty of decoding accuracy *r* in our analyses. Specifically, in the case of linear models, we *Z* -transform *r* relative to each decoder’s own empirically determined null distribution, thereby insulating against spurious correlations and facilitating comparison across features. We describe our approach in detail in the following sections.

## 2. Methods

Here, we test the neural decoding of an extended acoustic feature set drawn from psychophysics and music information retrieval research. The experimental data underlying the analyses in this study were also reported in [19]. In that analysis, we focused on methodological aspects of decoding the amplitude envelope only. In brief, the experiment consisted of an audiobook listening task and concurrent neural recording with EEG. Participants were presented with a complete Sherlock Holmes short story that was performed in English (intelligible; Duration = 33 min 47 sec) and Dutch (nonintelligible; Duration = 35 min 54 sec) by an early bilingual speaker. There were three acoustic conditions, consisting of natural speech, spectrally degraded but highly intelligible speech, and spectrally degraded speech with additional spectrotemporal distortions (“blurring”). This final condition is more challenging to understand, but is still intelligible on average (rated as Median 5.50, IQR = 2.00 on a scale 1 *−* 7 by the participants in this study). We describe these experimental conditions in more detail below. There were two trials for each of the 6 combinations of language and acoustic condition, the ordering of which was fully counterbalanced across participants. Story content was presented chronologically in both languages. Given that the effects of auditory attention on speech neural decoding are well documented [8, 37, 38], we developed an attentive behavioural task for both intelligible and unintelligible speech that involved detecting speech-based auditory targets in the form of short repeated phrases occurring every *∼* 45 s throughout the experiment. Performance was generally good with participants achieving a Mean Hit Rate of 0.89 (SD = 0.16) and a Mean False Alarm Rate of 0.09 (SD = 0.13) in the most difficult experimental condition, spectrally degraded Dutch with blurring. The entire experimental session, including preparation, lasted approximately 2 *−* 2.5 hours.

### 2.1. Participants

The sample included thirty-eight adults (22 female and 16 male; aged 18–35, Mean = 24.95, SD = 5.09) with self-reported typical hearing and no history of speech nor language-related disorders. Inclusion criteria were English spoken as a primary language and minimal experience with Dutch. The study received approval from the Cambridge Psychology Research Ethics Committee and was conducted in accordance with the Declaration of Helsinki. Participants were reimbursed for their participation and compensated for travel expenses.

### 2.2. Stimuli and Acoustic Feature Set

Speech stimuli recordings were sampled at 44.1 kHz and root mean square-normalized. To apply spectral degradation, we employed the SPIRAL vocoder [39]. This method simulates aspects of CI listening, particularly cross-channel interactions at the electrode-neural interface. Such interactions lead to distortions, or blurring, of spectral-temporal patterns in speech and increased difficulty in understanding [33, 40]. We generated what we term the *Vocoded* condition using 16 analysis filter bands without blurring, and the *Vocoded + Blurring* condition with 16 analysis filter bands and a cross-channel interaction decay slope of *−*16 dB/octave. These parameters were empirically found to approximate cochlear implant user speech reception scores in typically hearing listeners for various acoustic conditions [39, 41].

#### 2.2.1. Feature Extraction and Processing

To produce the baseline envelope feature (*Env*), we first passed the stimulus audio through an Equivalent Rectangular Bandwidth (ERB) filterbank with 32 filter analysis bands from 50 to 5000 Hz. The sub-band outputs were then rectified, subjected to power-law compression, and averaged to form the univariate envelope [42]. In addition, we calculated a second envelope-based feature, peaks in the first derivative of the envelope (*Env Deriv*). Sometimes called “acoustic edges”, this feature captures local maxima in the envelope rate of change–corresponding to vowel onsets in speech–and has been investigated in previous neural speech tracking research [43, 44]. To generate spectral features, we used the Timbre Toolbox for MATLAB (MathWorks, Natick, USA), a suite of audio analysis tools developed to model empirical data from perceptual studies and for computational tasks, such as musical instrument classification [27]. A selection of representative features are plotted alongside the speech spectrogram in Figure 1, Panel A. It should be noted that the feature profiles remain generally stable across experimental conditions (Figure 1, Panel B), meaning that any differences in decoding accuracy can be attributed to changes in the neural response rather than loss of information or injected noise as a result of the acoustic modifications. Features were again extracted from the output of the ERB filterbank. Parameters comprised a window length of 35 ms and hop size of 10 ms. To reduce the influence of noise and improve stability, we scaled the features by a heuristically determined signal intensity threshold with Gaussian smoothing. We initially computed all available spectral descriptors, but selected only a subset for further analysis following inspection of the correlation matrices (Figure 1, Panel C). A summary of the initial acoustic feature set is provided in Supplementary Material, Table A1 and readers are referred to [27] for full details concerning the formulas used to generate the spectral descriptors.

**Figure 1.**
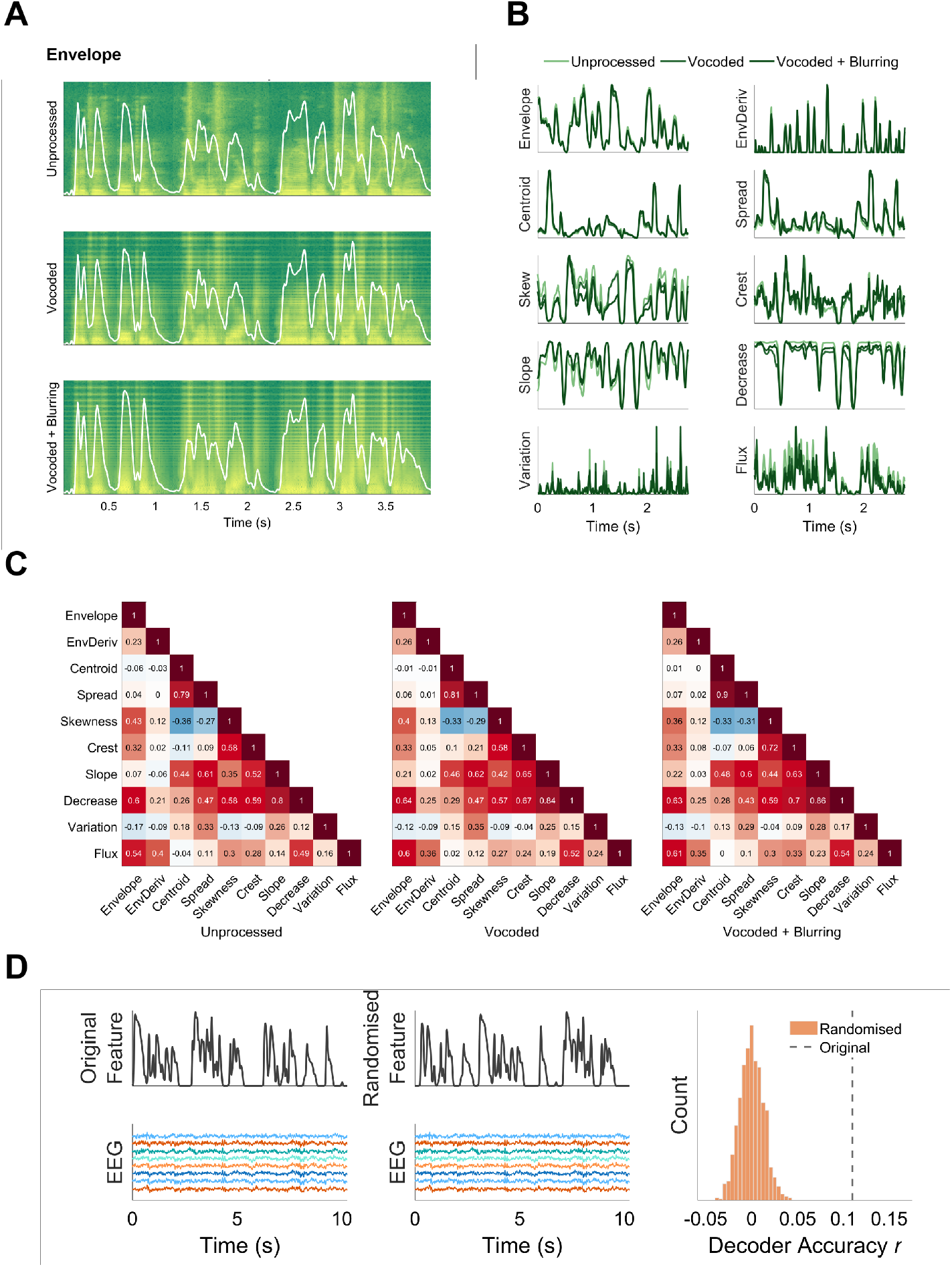
Panel A: Speech spectrograms for the three levels of spectral degradation with the corresponding envelope superimposed. Panel B: Acoustic features by level of spectral degradation. Panel C: Pearson’s correlation matrices depicting the relationships between acoustic features by level of spectral degradation. Panel D: Random permutation procedure used to generate null distributions for the linear decoding models.

### 2.3. Experimental Procedure

Following EEG cap preparation, participants were given a practice session with feedback automatically provided on detection and reaction times to auditory targets, which were measured using a handheld USB button device. After each trial presentation, they were invited to take a short, self-paced break. Participants were also asked to rate (1) how well they could understand and (2) how engaged they felt during the preceding story section on a seven-point Likert scale.

### 2.4. EEG Data Acquisition and Analysis

We recorded 64-channel EEG sampled at 2048 Hz using a BioSemi ActiveTwo EEG system (BioSemi, Amsterdam, Netherlands) with electrodes placed according to the International 10–20 system. Stimuli were presented via Etymotic Research (Elk Grove Village, USA) insert headphones in an electromagnetically shielded room. EEG data were re-referenced offline to the channel average and resampled at 256 Hz for computational efficiency. The data were highpass filtered at 0.5 Hz and then lowpass filtered at 40 Hz using fourth-order Butterworth filters. Eye blink artifacts were removed using independent component analysis and manually identified noisy channels were interpolated using a weighted neighbours approach. All preprocessing was performed in MATLAB [45] using Fieldtrip [46] and custom scripts. The first and final 5 s of each trial were excluded in case of filtering artifacts. Both the EEG and acoustic features were resampled to 100 Hz prior to decoder training.

#### 2.4.1. Decoding models

We trained individual linear decoders using regularised ridge regression as implemented by the Multivariate Temporal Function Toolbox in MATLAB [47]. A decoder reconstructs the stimulus feature as a linear convolution of the EEG over a range of time lags, which in this case was *−*50 to +500 ms post-stimulus onset. For each participant, acoustic feature, and experimental condition, we trained individual models with the regularisation parameter *λ* determined via *k* -fold cross-validation where *k* was 6 *−* 7, depending on the duration of training data. In most cases, *λ* was 100.

We also trained individual non-linear decoders using two types of deep neural networks (DNN): a fully-connected feed-forward neural network (FCNN) and a convolutional neural network (CNN) based on the EEGNet architecture [48]. We used similar methodology and network architectures as outlined in [5] for subject-specific decoding of the speech envelope from EEG recordings. The input data used for both DNNs spanned 500 ms (T=50 samples) across 64 recorded EEG channels (C=64 channels). Notably, the temporal receptive field is slightly wider in our study (500 ms) than in [5] (400 ms) due to a lower sampling rate of the EEG recordings (100 Hz). We trained both architectures to maximize the correlation between the predicted and ground truth speech features. For the FCNN, we fixed hyperparameters to match the population model in [5]: 3 hidden layers, dropout rate = 0.45, batch size = 256, and weight decay = 10^*−*4^. Similarly, for the CNN we used the population model hyperparameters: 8 filters (first convolutional layer), 64 filters (second and third layers), dropout rate = 0.20, batch size = 256, and weight decay = 10^*−*8^. For both architectures, the optimal learning rate was determined following a similar procedure as used in [5], by sampling from (10^*−*6^, 10^*−*5^, 10^*−*4^, 10^*−*3^, 10^*−*2^). Once the best learning rate was selected for a given participant, training was performed using this rate and automatically stopped after 30 epochs or if the correlation between speech features did not improve on the validation set for at least 5 consecutive epochs.

For all trained decoders, we obtained the dependent measure *r* by correlating the reconstructed and true input features using 60 s of held-out data selected from a mid-point in the experiment [4], with the remaining segments retained for training.

#### 2.4.2. Statistical analyses

Unless otherwise stated, dependent variables were analysed using linear mixed effect models using `lme4` [49] with post hoc tests conducted with the `emmeans` package [50]. Where relevant, model specification was guided using likelihood ratio tests and the Akaike information criterion (AIC). The significance of predictors was assessed using Type 2 Analysis of Variance tests with Satterthwaite’s degrees of freedom from the `lmerTest` package [51]. Random intercepts and slopes for participant are fit, subject to model convergence. Where appropriate, model reference (intercept) levels are set to Language: English, Spectral Degradation: Unprocessed, Feature: Envelope, and Model: Linear. Details concerning all reported models are provided in the Supplementary Material. For correlations, confidence intervals are generated via bootstrap with 5000 iterations.

#### 2.4.3. Random Permutations and Z-scoring of linear decoders

The range of decoding accuracy *r* may vary due to nuisance factors such as EEG system, sources of electrical and biological noise, and so forth. It is also possible that the decoding noise floor varies across different acoustic features, challenging their direct comparison. We, therefore, contextualise reconstructive accuracy *r* by generating a null distribution for each decoder (i.e., within participant, acoustic feature, and experimental condition) using randomised surrogate data (Figure 1, Panel D). Specifically, we algorithmically segmented each input feature at silent points into approximately 3 s chunks. These chunks were randomly permuted and re-concatenated using smoothed interpolation at join points. This process preserves local temporal structure, thereby ensuring the permuted feature is plausible, yet annihilating its long-term correspondence to the unmodified neural response. With each permutation, a new decoder is trained and used to reconstruct the intact held-out brain data. We run 1000 permutations per decoder to generate the dependent variable *Z*. Note that, due to the computationally demanding nature of this analysis, it was not possible to conduct random permutation testing with the non-linear models.

## 3. Results

### 3.1. Comparison of decoding performance across models

We first assessed decoding accuracy *r* across the three decoder architectures: Linear, FCNN, and CNN (Figure 2, Panel A). Linear mixed effects modelling (details in Supplementary Material, Table A2) confirmed that *r* significantly differed by model, *F* (2,6768) = 102.40, *p <* 0.001. Linear decoders produced the highest values on average (M = 0.12, SD = 0.07); followed by FCNN (M = 0.11, SD = 0.07), Est = *−*0.02, 95% CI [-0.03,-0.01], *t* (1,6768) = *−*4.30, *p <* 0.001; and then CNN (M = 0.10, SD = 0.08), Est = *−*0.03, 95% CI [-0.04,-0.02], *t* (1,6768) = *−*6.59, *p <* 0.001. In short, and contrary to previous reports [e.g., 5], we do not find an advantage for non-linear over linear decoders. There was also a significant main effect of feature, *F* (9,6768) = 479.26, *p <* 0.001, but no interaction between model and feature, *F* (18,6768) = 0.99, *p* = 0.462. This suggests the grouping of decoding accuracy *r* by feature is broadly consistent across architectures. We further quantify this result using Kendall’s Coefficient of Concordance *W*, a non-parametric measure of agreement where 0 indicates no agreement and 1 indicates perfect agreement [52]. We calculated *W* separately as *r* by participant ranked within experimental condition; and decoding accuracy *r* by experimental condition ranked within participant. High *W* would indicate that the decoders are sensitive to common inter- and within-subject neural signals, respectively. Mean *W* within experimental condition was 0.83 (SD = 0.07), indicating substantial agreement across decoders when ranking participants. The models also ranked experimental conditions *within* participant similarly, Mean *W* = 0.76 (SD = 0.16). This latter result is plotted by acoustic feature in Figure 2 (see also Supplementary Material, Figure A1 for bivariate analysis between models). In sum, we find a consistent pattern of decoding accuracy by feature and experimental condition, irrespective of model architecture.

**Figure 2.**
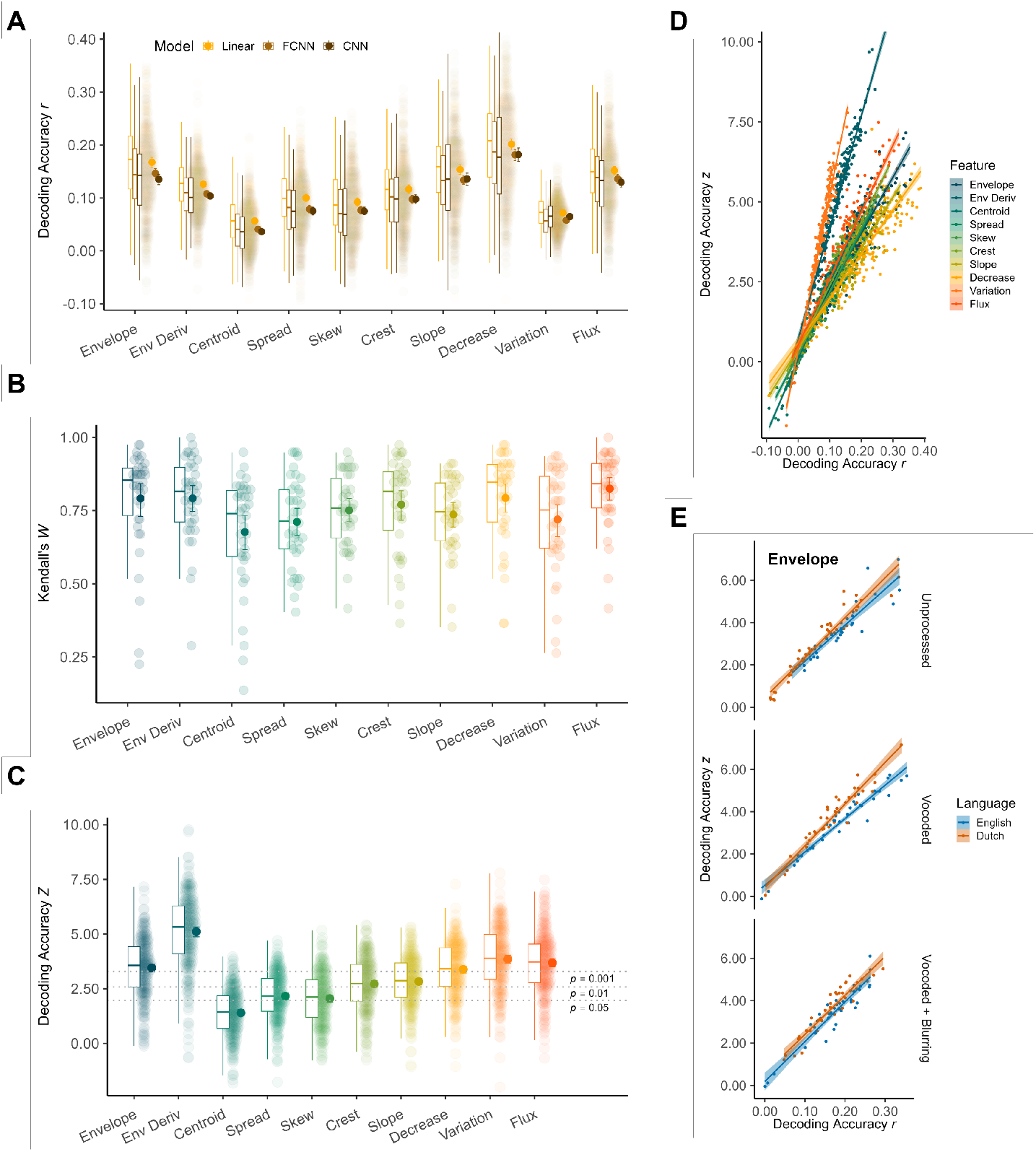
Panel A: Decoding accuracy (Pearson’s *r*) by acoustic feature and model architecture: Linear regression, fully connected neural network (FCNN) and convolutional neural network (CNN). Panel B: Agreement (Kendall’s Coefficient of Concordance *W*) of decoding accuracy (across experimental conditions and within participant), between model architectures by acoustic feature. Panel C: Decoding accuracy (*Z*) by feature with thresholds of statistical significance marked in dashed grey lines. Panel D: *Z* as a function of *r* by feature. Panel E: *Z* as a function of *r* by experimental condition within the *Envelope* feature. Box plots indicate the median and inter-quartile range of the distribution. Individual data points are pooled over participant and experimental condition. Solid markers represent the mean and error bars show the 95% confidence intervals of the mean. Shaded regions indicate 95% confidence intervals of the linear fit.

### 3.2. Assessing the robustness of individual acoustic feature decoding

Having established that linear and non-linear decoders yield comparable results, we turn now to the acoustic features themselves. We began by converting *r* to *Z* using randomly permuted data (see Section 2.4.3 for details). This procedure is motivated to compare the acoustic features on an equitable scale as well as providing an additional measure of robustness (i.e., as a *Z* -test). This step was infeasible for the computationally-intensive FCNN and CNN and so we focus here on linear decoders only. Averaging at the group level (Figure 2, Panel C), decoding accuracy surpasses *p <* 0.05 (*Z ≈* 1.96) for nearly every feature, with distributions of *Z* summarised in Supplementary Material, Table A3. From these data, we can reject the null hypothesis (i.e., that observed *r* has occurred by chance) for nine out of the ten acoustic features at *p* = 0.05, or five out of ten features at *p* = 0.001. When visually comparing *r* to *Z* (Panels A and C, Figure 2), it is apparent that their relationship is mediated by feature. In particular, *Spectral Variation* is associated with relatively low *r*, yet is the second-highest ranked feature by *Z*. We, therefore, examined the mapping of *r* to *Z* using a linear mixed effect model that included the interaction between *r* and feature, from which we could extract slopes (i.e., *Z ∼ r*; model details in Supplementary Material, Table A4). Averaging over experimental conditions, the coefficients by feature are reported in Supplementary Material, Table A5 and plotted in Figure 2, Panel D. Two relatively sparse features, *Spectral Variation* and *Env Deriv*, are distinguished by their steep slopes, but there is variability in general (see Supplementary Material, Table A6 for pairwise statistical comparisons). As an illustration, we can compare what values of *Z* are taken for different features when decoding accuracy *r* = 0.10: *Z* = 4.72 for *Spectral Variation, Z* = 2.03 for *Spectral Flux*, and *Z* = 1.34 for *Spectral Decrease*.

We also examined the relationship between *r* and *Z* across experimental conditions by running a separate mixed model within each feature. This exploratory approach yielded differences in the *Z ∼ r* slope according to experimental condition in three features: *Envelope, Env Deriv*, and *Spectral Spread*. For brevity, we report only the contrasts that reached corrected significance in Supplementary Material, Table A7, noting these were all for English versus Dutch within the same level of spectral degradation. For instance, Panel E in Figure 2 shows the difference between English and Dutch Vocoded speech when decoding *Envelope*, Estimate = 4.07, 95% CI [1.65, 6.50], *t* (214.00) = 3.31, *p*_*bonf*_ = 0.017, Cohen’s *d* = 0.45, 95% CI [0.18, 0.72]. Namely, the English slope (Coefficient = 15.38, 95% CI [13.80, 16.96]) is shallower than that of Dutch (Coefficient = 19.45, 95% CI [17.55, 21.35]), indicating a higher spurious correlation rate. This model suggests that achieving *Z* = 3.29 (*p* = 0.001) would require *r ≈* 0.18 for English and *r ≈* 0.15 for Dutch.

### 3.3. Sensitivity to spectral degradation and speech intelligibility

To test which features are most sensitive to the experimental manipulations, we performed a data-driven model selection procedure where we fit one predictor at a time for each feature. The final model specifications are reported in Table 1 with details and plots for all features in Supplementary Material Appendix A.6. For brevity, we report and visualise a representative subset, beginning with the simplest models, which contain only a random intercept. This group includes *Envelope* (Figure 3, Panel A). Although *Envelope* decoding accuracy is not affected by spectral degradation nor language, its conditional *R*^2^, which describes the total variance explained by the model, is 0.42. This indicates the relative stability of *Envelope* decoding within participant. There are also models where accuracy is mediated by spectral degradation only. *Spectral Slope* decoding, for instance, is enhanced with increasing spectral degradation, *F* (2.00,188.00) = 5.54, *p* = 0.005 (Figure 3, Panel B). Namely, Vocoded (Mean = 2.89, SD = 1.23) evokes significantly higher *Z* than Unprocessed (Mean = 2.55, SD = 0.95), Est = 0.34, 95% CI [0.03,0.65], *t* (1,188.00) = 2.18, *p* = 0.031. This is also the case for Vocoded + Blurring (Mean = 3.06, SD = 1.25), Est = 0.51, 95% CI [0.20,0.82], *t* (1,188.00) = 3.27, *p* = 0.001. But Vocoded and Vocoded + Blurring do not differ (*p*_*bonf*_ = 0.828).

**Table 1.** Final linear mixed effect models predicting decoding accuracy *Z* for each acoustic feature.

| Feature | Model | AIC | $R^2$ | |
| --- | --- | --- | --- | --- |
|  |  |  | Total | Fixed Effects |
| Envelope | Intercept | 742.14 | 0.42 | 0.00 |
| Env Deriv | Spectral Degradation | 794.96 | 0.64 | 0.01 |
| Centroid | Spectral Degradation | 691.47 | 0.19 | 0.04 |
| Spread | Intercept | 674.02 | 0.24 | 0.00 |
| Skew | Spectral Degradation + Language | 713.28 | 0.28 | 0.07 |
| Crest | Spectral Degradation + Language | 719.66 | 0.35 | 0.07 |
| Slope | Spectral Degradation | 691.46 | 0.33 | 0.03 |
| Decrease | Intercept | 715.21 | 0.39 | 0.00 |
| Variation | Spectral Degradation | 757.95 | 0.51 | 0.02 |
| Flux | Spectral Degradation + Language | 754.32 | 0.45 | 0.03 |

**Figure 3.**
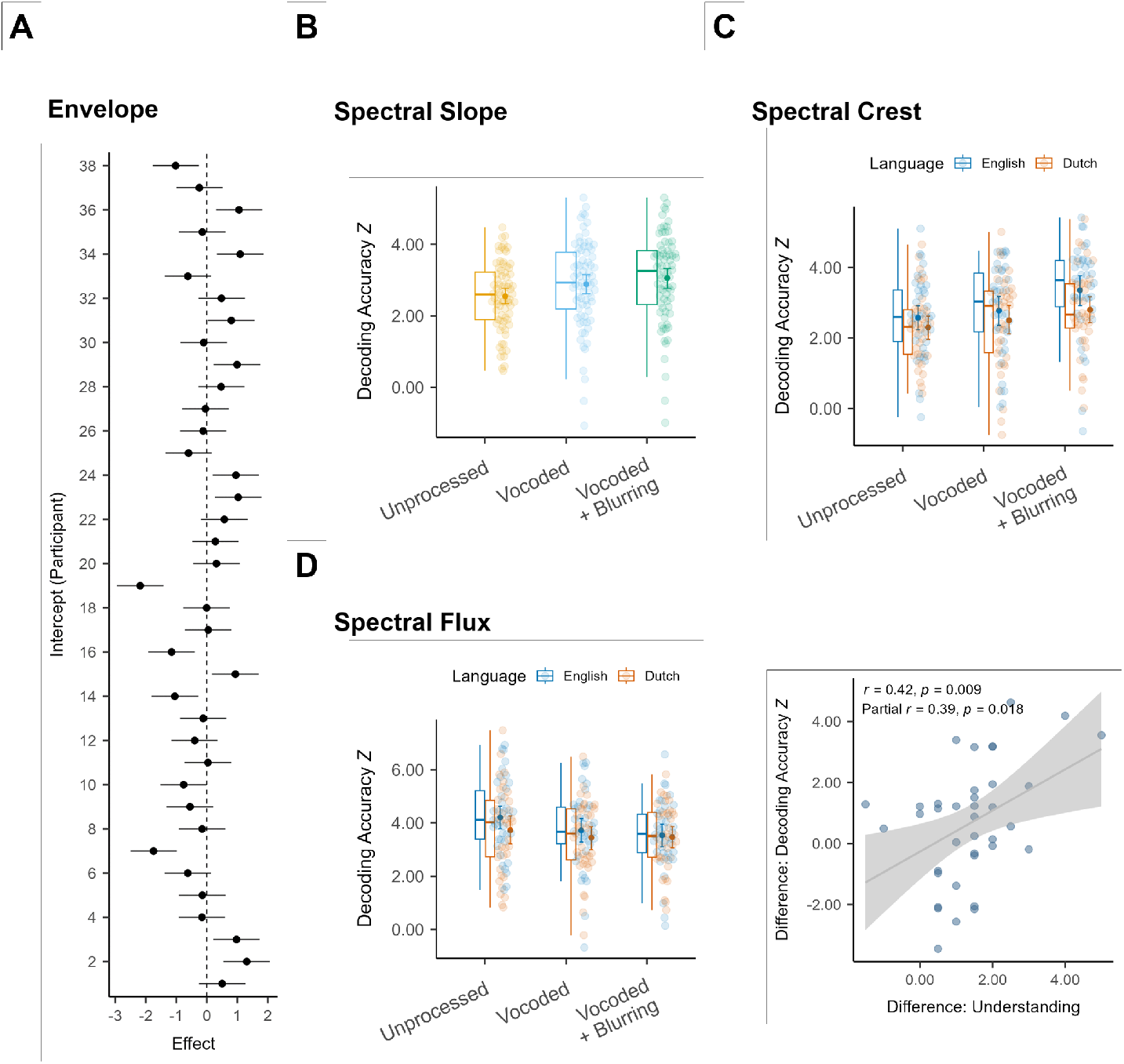
Panel A: Linear mixed effects model for *Envelope* decoding accuracy *Z*, which included only a random intercept for Participant. Markers and error bars indicate the intercept fit to each participant with 95% confidence intervals of the estimate. Panel B: Linear mixed effects model for *Spectral Slope* decoding accuracy *Z*, which included a fixed effect of Spectral Degradation and a random intercept for Participant. Panel C: Linear mixed effects model for *Spectral Crest* decoding accuracy *Z*, which included fixed effects of Spectral Degradation and Language and a random intercept for Participant. Panel D: Left, Linear mixed effects model for *Spectral Flux* decoding accuracy *Z*, which included a fixed effect of Spectral Degradation and a random intercept for Participant. Right, Pearson’s correlation and partial correlation between change in self-reported speech understanding between unprocessed and vocoded + blurring English, and change in decoding accuracy *Z* between unprocessed and vocoded + blurring English. Box plots indicate the median and inter-quartile range of the distribution. Individual data points are pooled over participant and experimental condition. Solid markers represent the mean and error bars show the 95% confidence intervals of the mean. Shaded regions indicate 95% confidence intervals of the linear fit.

Finally, the remaining models have an effect of both spectral degradation and language, but not their interaction. *Spectral Crest* decoding (Figure 3, Panel C) improves with increasing spectral degradation, *F* (2.00,187.00) = 7.95, *p <* 0.001. In particular, Vocoding + Blurring (Mean = 3.08, SD = 1.28) is associated with higher *Z* than both Unprocessed (Mean = 2.44, SD = 1.09), Est = 0.64, 95% CI [0.32,0.97], *t* (1,187.00) = 3.90, *p <* 0.001) and Vocoded (Mean = 2.64, SD = 1.31), Est = 0.44, 95% CI [0.11,0.76], *t* (1,222.00) = 2.67, *p*_*bonf*_ = 0.025. Unprocessed and Vocoded, however, do not differ (*p* = 0.219). In addition, the effect of language on *Crest* decoding is also significant, *F* (2.00,187.00) = 7.95, *p <* 0.001, with English (Mean = 2.90, SD = 1.29) more accurate than Dutch (Mean = 2.53, SD = 1.19), Est = 2.62, 95% CI [2.28,2.97], *t* (1,108.03) = 15.10, *p <* 0.001. To highlight one last feature, we turn to *Spectral Flux* (Figure 3, Panel D, left). Unlike with *Slope* and *Crest*, the effect of spectral degradation was significantly negative, *F* (2.00,187.00) = 4.20, *p* = 0.016. Unprocessed (Mean = 3.97, SD = 1.53) was decoded more accurately than Vocoded (Mean = 3.58, SD = 1.42), Est = *−*0.39, 95% CI [-0.72,-0.05], *t* (1,187.00) = *−*2.25, *p* = 0.026, as well as Vocoded + Blurring (Mean = 3.50, SD = 1.26), Est = *−*0.47, 95% CI [-0.80,-0.13], *t* (1,187.00) = *−*2.71, *p* = 0.007. Vocoded and Vocoded + Blurring did not differ from one another (*p*_*bonf*_ = 1.00). There was also a marginally-significant effect of language, Est = *−*0.28, 95% CI [-0.55,0.00], *t* (1,187.00) = *−*1.96, *p* = 0.051. As with *Crest*, English (Mean = 3.82, SD = 1.42) was decoded slightly better than Dutch (Mean = 3.55, SD = 1.41). We note that the features with higher *Z* on average also have larger conditional *R*^2^–in fact, the variables are nearly perfectly correlated (rho = 0.98). Features with conditional *R*^2^ *>* 0.40 also have relatively low marginal *R*^2^ *≤* 0.03. Together, this may suggest a trade-off between the detection by EEG of stable, generalised auditory processing, versus neural markers that are noisier, but more contextually sensitive to speech understanding and/or spectral fine detail.

### 3.4. Correlation with self-reported understanding

For English speech, participants rated the unprocessed and vocoded acoustic conditions at ceiling level for intelligibility on average. Although vocoded + blurring received a median rating of 5.5 out of 7 for self-reported understanding, individual listeners varied in how well they felt they could follow the content of the story. In our previous analysis [19], we observed a Pearson correlation (denoted here as *R*) between changes in envelope decoding and changes in self-reported understanding for English. Namely, participants whose decoding accuracy *r* dropped from unprocessed to vocoded + blurring also tended to report a greater loss in understanding, *R* = 0.35, 95% CI [0.02, 0.60], *p* = 0.030. When we partialled out the equivalent neural change for Dutch speech, however, the correlation diminished (*R* = 0.29, 95% CI [*−*0.11, 0.60], *p* = 0.077), suggesting the relationship was partly driven by acoustic changes rather than loss in intelligibility *per se*. Here, we sought to confirm our original result in the *Z* domain, and gauge whether other acoustic features may track changes in speech understanding more closely. With respect to *Envelope, R* = 0.35, 95% CI [*−*0.01, 0.66], *p* = 0.031–essentially the same result as for the decoding accuracy *r*. Across the other features, most had *R ≤* 0.30 and uncorrected *p ≥* 0.070. The only exception was for *Spectral Flux* (Figure 3, Panel D, right). Without accounting for acoustic differences, *R* = 0.42, 95% CI [0.14, 0.62], *p* = 0.009. When we introduce the covariate, Dutch unprocessed to vocoded + blurring neural difference, then *R* = 0.39, 95% CI [0.14, 0.58], *p* = 0.018. Hence, the *Spectral Flux* correlation is mostly preserved when controlling for acoustically-driven changes in decoding accuracy. Yet, while they do not include 0, the wide confidence intervals of the estimate indicate its uncertainty.

## 4. Discussion

The current study assesses the neural decoding of a set of acoustic features. Going beyond the amplitude envelope and its derivative, we quantified the robustness of spectral descriptors that convey how the frequency characteristics of speech sounds are shaped and change over time. This robustness was established partly by employing three independent linear and non-linear model architectures, and partly by conducting an extensive permutation analysis of the linear decoders. Finally, we examined how feature decoding varies according to acoustic clarity and speech understanding. Our approach shows that, whereas the neural decoding of some features appears to reflect generalised–and likely low-level–auditory processing, others may shed light on which spectral properties are more relevant during speech perception and comprehension. In the following section, we discuss our findings with their gains and limitations, and conclude with a view to potential applications.

### 4.1. Comparable decoding across linear and DNN decoders

Employing three contrasting architectures, we found comparable decoding accuracy at the model level, with slightly higher decoding accuracy *r* associated with linear regression-based decoders but comparable variability. Furthermore, the patterning of accuracy by feature and experimental condition was broadly similar, despite the theoretical advantages conferred by non-linear decoders. That is to say, we foresee no obvious risk of information loss by choosing the computationally less demanding linear model, at least with respect to our experimental manipulations and feature set. Moreover, the convergences across architectures suggest that the neural tracking of speech acoustics can be operationalised as a linear transformation.

This conclusion differs from [5], wherein the authors report clear benefits to decoding with the two DNN approaches. Although we directly applied the publicly available code and analysis pipeline from that study for subject-specific training of decoders, there were some important methodological differences. In particular, we fixed the model hyperparameters *a priori* that were individually optimised in [5], and only optimised the learning rate on a subject-by-subject basis. In our view, this approach may be more analogous to optimising the regularisation hyperparameter of the linear regression models, enabling us to cross-check feature decoding across architectures and perform a fair comparison. It is likely that tuning many hyperparameters via random search would boost individual DNN decoding, but it limits generalisation of the approach and requires more computational resources. Given the recurring pattern of results by feature and experimental condition across decoders, we have some assurance that the decoders did not overfit and their predictions agree in reflecting the underlying pattern in the data. Finally, we find that accuracy *r* does not translate to *Z* in a uniform manner across features. Hence, a higher or lower DNN decoding accuracy *r* may not necessarily be more or less informative or reliable per se.

### 4.2. How best to estimate decoding accuracy?

It is recommended to compare decoding accuracy *r* with a null distribution in order to determine decoder significance (i.e., where observed *r* exceeds *n*^*th*^ null percentile) [4]. Yet, in the literature, group-level comparisons are typically conducted upon the *r* values themselves. Our analysis revealed differences in the *r* to *Z* mapping that are sizeable enough to impact binary decisions regarding statistical significance within a range of *r* typical of the literature. This disparity occurred across features, as well as within. In particular, we saw that, for the envelope, English elicited higher *r* values than Dutch speech by chance. As many speech decoding studies contrast intelligible and non-intelligible languages, this result underscores the potential consequence of interpreting *r* directly without empirically estimating its uncertainty. Hence, rather than make *r* the object of analysis with significance determined as an arbitrary cut-off, we propose *Z* as a more reliable outcome variable–or at least, reporting both. *Z* is continuous and integrates important information about sampling uncertainty. Moreover, rescaling decoding accuracy within *Z* -space may help reconcile results across study sites or account for other nuisance differences, such as feature sparsity–noting that *Z* will also depend on the number of permutations performed and the permutation method itself. One can argue that statistical significance is different from magnitude of the effect. Acknowledging this, we counter that the meaning and interpretation of *r* remains an open question [3], especially when *r* may be subject to inflation, as we have observed in the present data. Finally, we note that *Z*, at least, provides an intuitive and empirically determined measure of the distance between observed *r* and 0, which is the mean of a distribution of *r* where decoding has occurred entirely coincidentally. Thus, although more work is needed to understand the specific neural computations that give rise to a given feature’s decodability, *Z* quantifies the statistical robustness of any observed stimulus-brain correspondence and can be a powerful tool for making comparisons where there is a risk of different noise floors (for example across studies, participants or acoustic features).

### 4.3. Robust auditory decoding and sensitivity to experimental manipulations

According to the random permutation analysis, the most robustly decoded acoustic features were *Env Deriv, Spectral Variation, Spectral Flux*, and *Envelope*. These features were decoded similarly well for intelligible and non-intelligible speech, with the exception of a marginal effect for *Spectral Flux* only. In the current study, we also noticed that *Z* has a close association with conditional *R*^2^, which primarily reflects variance explained by participant. This means that, for features with higher *Z* on average, *Z* is also more stably distributed within participants. What this suggests to us is that differences in intelligibility, which we did observe for other features, may reflect a loss of decodability for incomprehensible speech, rather than a gain for speech that is understood. Thus, although our results identify candidate acoustic features that may differentiate understanding, their use for this purpose would come at the cost of an overall noisier and less reliable neural signal. This also holds for features that may discriminate between levels of acoustic clarity, such as *Spectral Slope*.

With respect to *Envelope*, the null effect of intelligibility in the *Z* domain is at odds with our findings in [19], where we observed a main effect of language on envelope decoding measured as *r*. But the disparity can be explained by the aforementioned language-based differences in the linear relationship between *r* and *Z*. In contrast, we do see small but significant effects of language for *Spectral Crest* and *Spectral Skew*. These features are moderately correlated with each other and probably map onto a common linguistic component. For instance, *Spectral Crest* describes how peaked the spectrum is, roughly approximating the tonal-to-noise ratio of the sound. In speech, this would likely correspond to vowels and voiced segments. When speech is understood, more neural resources could be allocated to the online analysis of these components, resulting in the increased decoding accuracy for English we observed. Conversely, whereas vocoding speech removes fine spectral detail, it also concentrates spectral energy within wide, defined bands. This may explain why decoding accuracy also tends to increase with spectral degradation for *Spectral Crest* and *Spectral Skew*. By comparison, decoding of *Spectral Flux*, a measure of spectral change theorised to correspond to phoneme onsets [36], was deleteriously affected by vocoding. It is, therefore, possible that processing of *Flux* operates on more fine-grain spectral information than *Spectral Skew* and *Spectral Crest*. We note that individual variability in *Spectral Flux* decoding was uniquely correlated to self-reported difficulties understanding English under spectral degradation, showing potential promise–and warranting further scrutiny–for future studies.

### 4.4. Limitations and clinical outlook

This study has several important limitations. With respect to model architecture, as discussed above, we opted not to tune multiple DNN hyperparameters for individual listeners when training subject-specific decoders. This allowed us to generalise across participants and features, but our results may not fully reflect the upper limits of non-linear model performance. We were also precluded from performing the permutation analyses on the DNN approaches, given the time and computational resources required to re-train the models hundreds of times. An alternative choice would be a compromise by using the non-randomised DNN model to predict mismatched EEG and speech segments [5]. However, this is not a pure test of the decoder *per se* and the underlying hypothesis is different. Given we found strong agreement with respect to participant, feature, and experimental condition ranking across the linear and non-linear models, we opted to move forward with the linear decoders only and generate null distributions by retraining the decoder and applying it to matched segments (i.e., we are asking how well a chance-trained decoder can detect a truly present relationship).

Another caveat is with respect to the features themselves. We selected low level features originally developed for use in psychophysics and musical timbre research, wherein short sound stimuli are often synthesised and tightly controlled. Continuous, naturally produced speech is likely to produce noisier estimates, although we took steps in our methods to address this. Moreover, the relationship between low level acoustic descriptors and phonetic or other linguistically defined units is unclear and an active area of empirical inquiry [21, 53]. Thus, we can only speculate that, for instance, *Spectral Crest* may be related to voiced segments. To this point, it is possible that linguistic feature decoding–e.g., of word onsets–would correlate more strongly with comprehension (as discussed by [54, 55] but see also [56]). However, the preparation of these features requires expert annotation and/or the correction of forced alignment procedures. Moreover, phonetic or lexical decoding may not be useful in the context of prelingual infants or where verbal communication is otherwise impossible. In that case, evaluating the transmission of acoustic features–which could yet be relevant for future language acquisition or rehabilitation–would be paramount. Hence, although they are inherently noisy and difficult to directly assimilate into existing phonetic theory, acoustic features offer other advantages: They are defined by the physical signal (i.e., unlike phonemes), straightforwardly interpretable (unlike machine learning-derived features or cepstral coefficients–though see [53]), and easy to produce (unlike annotations).

One hope for speech decoding is that it could potentially index the fidelity with which assistive hearing devices, such as CI, transmit sound signals to the cortex and help in the fitting and optimisation of device parameters. Although we observed modest effects of spectral degradation, as well as intelligibility, they are too noisy to be used to make assessments on an individual level. We, therefore, close by considering two outlooks for future research into clinical applications. The first is that, on the one hand, the EEG signal to noise ratio might simply be too poor for obtaining fine-grained audiological insights. On the other hand, a parametric approach where the stimulus is altered on a continuum could provide an individualised estimate, for example, of spectral resolution by finding the kneepoint at which decoding accuracy *Z* drops to approximately zero [40, 33]. This relativistic approach may help with overcoming the large uncertainty that accompanies absolute decoding accuracy measures. The other direction to explore is EEG *encoding* analysis, which is a closely related method for estimating speech-brain coupling. Unlike decoders, encoding models are causal and the weights can be interpreted as spatio-temporal patterns of activation in sensor space [57, 2]. We found in this data set that the most robustly decoded features were less affected by experimental manipulations, suggesting their successful decoding was strongly driven by early auditory processing. Thus, an encoder-oriented analysis could enable the investigation of more focal effects that are essentially swamped by a relatively sizeable acoustic response.

## 5. Data availability statement

The data and scripts that support the findings of this study will be made openly available in a dedicated repository (OSF - URL forthcoming). For the purpose of open access, the authors have applied a Creative Commons attribution (CC BY) license to any author accepted manuscript version arising.

## 7. Authors’ contributions

A.D.M.: Conceptualization, Methodology, Programming, Formal analysis, Investigation, Writing - Original Draft, Visualization, Funding acquisition; C.G.: Methodology, Software, Formal analysis, Writing - Original Draft; TG: Conceptualization, Programming, Formal analysis, Investigation, Writing - Original Draft, Funding acquisition.

## 8. Acknowledgements

A.D.M. was supported by a Leverhulme Trust (ECF-2023-539) and Isaac Newton Trust Early Career Fellowship Award. C.G. was supported by a Pasteur-Roux-Cantarini Fellowship of Institut Pasteur, fundings from the Fondation Pour l’Audition (Grant No. FPA RD-2021-1, FPA IDA10), and a French government grant managed by the Agence Nationale de la Recherche under the France 2030 program, (ANR-23-IAHU-0003). T.G. was funded by Medical Research Council UK (Grant No. MR/T03095X/1).

## Appendix A. Supplementary Material

### Appendix A.1. Acoustic Feature Selection

**Table A1.** Summary of the acoustic feature set. Redundant highly correlated features (i.e., Mean *r ≥* 0.85) are removed. Full details concerning the generation of the spectral descriptors are provided in [27].

| Feature | Description | Retained | $r$ |
| --- | --- | --- | --- |
| *Env* | Amplitude envelope as calculated in X | Yes |  |
| *Env Deriv* | Peaks in the envelope first derivative | Yes |  |
| *Spectral Centroid* | Amplitude-weighted mean of the frequency representation | Yes |  |
| *Spectral Spread* | Standard deviation of the spectrum around its centroid | Yes |  |
| *Spectral Skew* | Asymmetry of the spectrum around its centroid | Yes |  |
| *Spectral Kurtosis* | Relative weight of the spectral distribution tails | No | Skew $\geq 0.95$ |
| *Spectral Flatness* | Ratio of the spectral geometrical mean to its arithmetic mean (approximates noise-to-tonality) | No | Spread $\geq 0.85$ |
| *Spectral Crest* | Ratio of the spectral maximum to its arithmetic mean (approximates noise-to-tonality) | Yes |  |
| *Spectral Slope* | Gradation of the spectrum envelope as determined by linear regression over its amplitude values | Yes |  |
| *Spectral Decrease* | An alternative calculation of spectral slope that emphasises lower frequencies | Yes |  |
| *Spectral Roll Off* | The frequency below which 95% of signal energy is contained | No | Centroid $\geq 0.87$ |
| *Spectral Flux* | Spectral change expressed as a difference over adjacent windows | Yes |  |
| *Spectral Variation* | Spectral change expressed as a normalised correlation over adjacent windows | Yes |  |

### Appendix A.2. Comparison of decoding performance across model architectures

**Figure A1.**
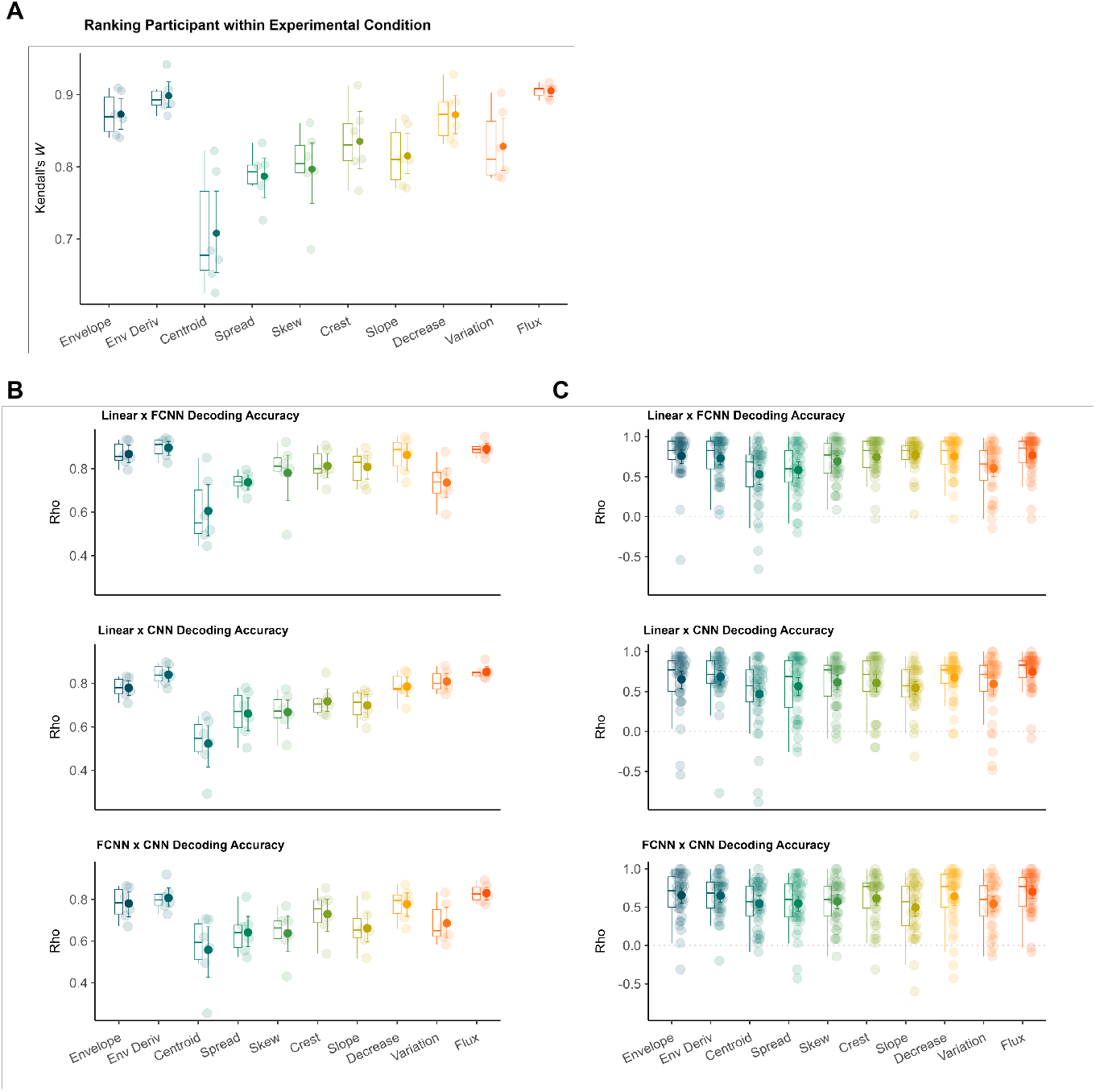
Panel A: Agreement (Kendall’s Coefficient of Concordance *W*) of decoding accuracy (within experimental condition and across participants) between model architectures by acoustic feature. Panels B and C: Bivariate association (Spearman’s rho) of decoding accuracy by acoustic feature and model architecture: Linear regression, fully connected neural network (FCNN) and convolutional neural network (CNN). Panel B shows correlations within experimental condition and across participants, and Panel C shows correlations within participant and across experimental conditions. Box plots indicate the median and inter-quartile range of the distribution. Individual data points are pooled over participant and experimental condition. Solid markers represent the mean and error bars show the 95% confidence intervals of the mean.

### Appendix A.3. Comparison of linear, FCNN, and CNN models

**Table A2.** Model formula: *r ∼* Model *×* Feature + Spectral Degradation *×* Language + (1 | Participant). Standardized parameters were obtained by fitting the model on a standardized version of the dataset. 95% Confidence Intervals (CIs) and p-values were computed using a Wald t-distribution approximation.

| Parameter | Estimate | 95% CI | | $t$ | df | $p$ | Std Est | 95% CI | |
| --- | --- | --- | --- | --- | --- | --- | --- | --- | --- |
|  |  | Lower | Upper |  |  |  |  | Lower | Upper |
| (Intercept) | 0.17 | 0.16 | 0.18 | 26.23 | 79.57 | < 0.001 | 0.80 | 0.63 | 0.98 |
| FCNN | -0.02 | -0.03 | -0.01 | -4.30 | 6768.00 | < 0.001 | -0.28 | -0.41 | -0.15 |
| CNN | -0.03 | -0.04 | -0.02 | -6.59 | 6768.00 | < 0.001 | -0.43 | -0.56 | -0.30 |
| Env Deriv | -0.04 | -0.05 | -0.03 | -8.43 | 6768.00 | < 0.001 | -0.55 | -0.68 | -0.42 |
| Centroid | -0.11 | -0.12 | -0.10 | -22.61 | 6768.00 | < 0.001 | -1.47 | -1.60 | -1.34 |
| Spread | -0.07 | -0.08 | -0.06 | -13.72 | 6768.00 | < 0.001 | -0.89 | -1.02 | -0.76 |
| Skew | -0.08 | -0.08 | -0.07 | -15.30 | 6768.00 | < 0.001 | -0.99 | -1.12 | -0.87 |
| Crest | -0.05 | -0.06 | -0.04 | -10.38 | 6768.00 | < 0.001 | -0.67 | -0.80 | -0.55 |
| Slope | -0.01 | -0.02 | 0.00 | -2.85 | 6768.00 | 0.004 | -0.18 | -0.31 | -0.06 |
| Decrease | 0.03 | 0.02 | 0.04 | 6.92 | 6768.00 | < 0.001 | 0.45 | 0.32 | 0.58 |
| Variation | -0.10 | -0.11 | -0.09 | -19.46 | 6768.00 | < 0.001 | -1.26 | -1.39 | -1.14 |
| Flux | -0.02 | -0.03 | -0.01 | -3.18 | 6768.00 | 0.001 | -0.21 | -0.33 | -0.08 |
| Vocoded | 0.00 | 0.00 | 0.01 | 2.03 | 6768.00 | 0.042 | 0.06 | 0.00 | 0.12 |
| Vocoded + Blurring | 0.01 | 0.00 | 0.01 | 2.49 | 6768.00 | 0.013 | 0.07 | 0.02 | 0.13 |
| Language | -0.02 | -0.03 | -0.02 | -10.34 | 6768.00 | < 0.001 | -0.30 | -0.36 | -0.24 |
| FCNN×Env Deriv | 0.00 | -0.01 | 0.02 | 0.42 | 6768.00 | 0.674 | 0.04 | -0.14 | 0.22 |
| CNN×Env Deriv | 0.01 | 0.00 | 0.02 | 1.43 | 6768.00 | 0.152 | 0.13 | -0.05 | 0.31 |
| FCNN×Centroid | 0.01 | -0.01 | 0.02 | 0.79 | 6768.00 | 0.429 | 0.07 | -0.11 | 0.25 |
| CNN×Centroid | 0.01 | 0.00 | 0.03 | 1.77 | 6768.00 | 0.077 | 0.16 | -0.02 | 0.34 |
| FCNN×Spread | 0.00 | -0.01 | 0.01 | -0.03 | 6768.00 | 0.980 | 0.00 | -0.18 | 0.18 |
| CNN×Spread | 0.01 | -0.01 | 0.02 | 1.10 | 6768.00 | 0.270 | 0.10 | -0.08 | 0.28 |
| FCNN×Skew | 0.01 | -0.01 | 0.02 | 0.82 | 6768.00 | 0.411 | 0.08 | -0.10 | 0.26 |
| CNN×Skew | 0.02 | 0.00 | 0.03 | 2.17 | 6768.00 | 0.030 | 0.20 | 0.02 | 0.38 |
| FCNN×Crest | 0.00 | -0.01 | 0.02 | 0.30 | 6768.00 | 0.768 | 0.03 | -0.15 | 0.21 |
| CNN×Crest | 0.01 | 0.00 | 0.03 | 1.96 | 6768.00 | 0.050 | 0.18 | 0.00 | 0.36 |
| FCNN×Slope | 0.00 | -0.01 | 0.01 | 0.19 | 6768.00 | 0.852 | 0.02 | -0.16 | 0.20 |
| CNN×Slope | 0.01 | 0.00 | 0.03 | 2.11 | 6768.00 | 0.035 | 0.19 | 0.01 | 0.37 |
| FCNN×Decrease | 0.00 | -0.01 | 0.01 | 0.18 | 6768.00 | 0.858 | 0.02 | -0.16 | 0.20 |
| CNN×Decrease | 0.01 | 0.00 | 0.03 | 1.85 | 6768.00 | 0.064 | 0.17 | -0.01 | 0.35 |
| FCNN×Variation | 0.01 | -0.01 | 0.02 | 0.99 | 6768.00 | 0.321 | 0.09 | -0.09 | 0.27 |
| CNN×Variation | 0.03 | 0.01 | 0.04 | 3.62 | 6768.00 | < 0.001 | 0.33 | 0.15 | 0.51 |
| FCNN×Flux | 0.01 | -0.01 | 0.02 | 0.74 | 6768.00 | 0.461 | 0.07 | -0.11 | 0.25 |
| CNN×Flux | 0.01 | 0.00 | 0.02 | 1.45 | 6768.00 | 0.146 | 0.13 | -0.05 | 0.31 |
| Vocoded×Language | 0.01 | 0.00 | 0.02 | 2.91 | 6768.00 | 0.004 | 0.12 | 0.04 | 0.20 |
| Vocoded + Blurring×Language | 0.01 | 0.01 | 0.02 | 4.31 | 6768.00 | < 0.001 | 0.18 | 0.10 | 0.26 |
| Random Effects | Variance | SD | ICC | Model Metrics |  |  |  |  |  |
| Within-Group Variance | 0.003 | (0.052) |  | AIC |  |  |  | -20380.54 |  |
| Between-Group Variance |  |  |  | BIC |  |  |  | -20127.81 |  |
| Random Intercept (Participant) | 0.001 | (0.033) | 0.534 | R<sup>2</sup> (Total) |  |  |  | 0.52 |  |
|  |  |  |  | R<sup>2</sup> (Fixed Effects) |  |  |  | 0.33 |  |
| N Groups |  |  |  | ICC |  |  |  | 0.28 |  |
| Participant | 38 |  |  | RMSE |  |  |  | 0.05 |  |
| N Observations | 6840 |  |  |  |  |  |  |  |  |

### Appendix A.4. Summary of Z

**Table A3.** Descriptive statistics summarising *Z* by acoustic feature.

*Appendix A.4. Summary of $Z$* **Table A3.** Descriptive statistics summarising $Z$ by acoustic feature.
| Feature | Mean | SD | 95% CI |  | Quantiles |  |  |  |  |
| --- | --- | --- | --- | --- | --- | --- | --- | --- | --- |
|  |  |  | Lower | Upper | 5<sup>th</sup> | 25<sup>th</sup> | 50<sup>th</sup> | 75<sup>th</sup> | 95<sup>th</sup> |
| Env Deriv | 5.11 | 1.81 | 4.87 | 5.35 | 1.94 | 4.10 | 5.32 | 6.29 | 7.42 |
| Variation | 3.85 | 1.49 | 3.65 | 4.04 | 1.41 | 2.92 | 3.89 | 4.98 | 6.04 |
| Flux | 3.69 | 1.42 | 3.50 | 3.87 | 1.13 | 2.77 | 3.73 | 4.54 | 5.99 |
| Envelope | 3.45 | 1.38 | 3.27 | 3.63 | 0.83 | 2.58 | 3.56 | 4.43 | 5.50 |
| Decrease | 3.38 | 1.28 | 3.21 | 3.55 | 1.17 | 2.60 | 3.41 | 4.38 | 5.14 |
| Slope | 2.83 | 1.17 | 2.68 | 2.98 | 0.86 | 2.11 | 2.87 | 3.68 | 4.58 |
| Crest | 2.72 | 1.25 | 2.55 | 2.88 | 0.45 | 1.93 | 2.73 | 3.61 | 4.47 |
| Spread | 2.16 | 1.09 | 2.02 | 2.31 | 0.31 | 1.47 | 2.17 | 2.98 | 3.78 |
| Skew | 2.05 | 1.20 | 1.89 | 2.21 | 0.06 | 1.19 | 2.12 | 2.91 | 3.88 |
| Centroid | 1.40 | 1.11 | 1.25 | 1.54 | -0.45 | 0.69 | 1.44 | 2.18 | 3.07 |

### Appendix A.5. Z as a function of r by Feature and Experimental Condition

**Table A4.** Model formula: *Z ∼ r ×* Feature + Spectral Degradation *×* Language + (1 | Participant). Standardized parameters were obtained by fitting the model on a standardized version of the dataset. 95% Confidence Intervals (CIs) and p-values were computed using a Wald t-distribution approximation.

| Parameter | Estimate | 95% CI |  | *t* | df | *p* | Std Est | 95% CI |  |
| --- | --- | --- | --- | --- | --- | --- | --- | --- | --- |
|  |  | Lower | Upper |  |  |  |  | Lower | Upper |
| (Intercept) | 0.57 | 0.43 | 0.71 | 8.10 | 1609.60 | < 0.001 | -0.23 | -0.28 | -0.19 |
| r | 16.97 | 16.27 | 17.67 | 47.41 | 2253.04 | < 0.001 | 0.75 | 0.72 | 0.79 |
| Env Deriv | 0.24 | 0.05 | 0.42 | 2.47 | 2225.74 | 0.013 | 1.39 | 1.34 | 1.43 |
| Centroid | -0.50 | -0.65 | -0.35 | -6.58 | 2233.67 | < 0.001 | 0.13 | 0.07 | 0.19 |
| Spread | -0.31 | -0.48 | -0.15 | -3.72 | 2224.43 | < 0.001 | -0.06 | -0.11 | -0.02 |
| Skew | -0.35 | -0.51 | -0.20 | -4.42 | 2225.80 | < 0.001 | -0.03 | -0.08 | 0.02 |
| Crest | -0.17 | -0.34 | 0.00 | -1.99 | 2221.56 | 0.047 | 0.09 | 0.04 | 0.14 |
| Slope | -0.15 | -0.33 | 0.02 | -1.72 | 2221.11 | 0.086 | -0.20 | -0.25 | -0.16 |
| Decrease | 0.07 | -0.11 | 0.25 | 0.72 | 2217.56 | 0.471 | -0.22 | -0.28 | -0.17 |
| Variation | -0.16 | -0.34 | 0.03 | -1.69 | 2225.64 | 0.091 | 2.14 | 2.07 | 2.21 |
| Flux | 0.00 | -0.18 | 0.18 | -0.01 | 2217.17 | 0.994 | 0.24 | 0.19 | 0.29 |
| Vocoded | -0.04 | -0.10 | 0.01 | -1.49 | 2215.94 | 0.136 | -0.03 | -0.06 | 0.01 |
| Vocoded + Blurring | 0.01 | -0.05 | 0.06 | 0.33 | 2216.20 | 0.738 | 0.01 | -0.03 | 0.04 |
| Language | 0.07 | 0.01 | 0.12 | 2.42 | 2220.74 | 0.016 | 0.04 | 0.01 | 0.07 |
| r×Env Deriv | 16.83 | 15.59 | 18.06 | 26.73 | 2232.01 | < 0.001 | 0.75 | 0.69 | 0.80 |
| r×Centroid | 5.81 | 4.52 | 7.10 | 8.82 | 2224.26 | < 0.001 | 0.26 | 0.20 | 0.32 |
| r×Spread | 1.66 | 0.50 | 2.81 | 2.81 | 2222.42 | 0.005 | 0.07 | 0.02 | 0.13 |
| r×Skew | 2.47 | 1.36 | 3.57 | 4.39 | 2221.62 | < 0.001 | 0.11 | 0.06 | 0.16 |
| r×Crest | 2.56 | 1.48 | 3.65 | 4.64 | 2221.23 | < 0.001 | 0.11 | 0.07 | 0.16 |
| r×Slope | -1.51 | -2.51 | -0.51 | -2.97 | 2221.80 | 0.003 | -0.07 | -0.11 | -0.02 |
| r×Decrease | -3.56 | -4.47 | -2.65 | -7.67 | 2218.41 | < 0.001 | -0.16 | -0.20 | -0.12 |
| r×Variation | 30.21 | 28.36 | 32.07 | 31.94 | 2240.42 | < 0.001 | 1.34 | 1.26 | 1.43 |
| r×Flux | 3.28 | 2.23 | 4.32 | 6.16 | 2217.97 | < 0.001 | 0.15 | 0.10 | 0.19 |
| Vocoded×Language | 0.09 | 0.02 | 0.17 | 2.36 | 2216.30 | 0.018 | 0.06 | 0.01 | 0.10 |
| Vocoded + Blurring×Language | -0.01 | -0.09 | 0.07 | -0.19 | 2217.11 | 0.850 | 0.00 | -0.05 | 0.04 |
| Random Effects | Variance | SD | ICC | Model Metrics |  |  |  |  |  |
| Within-Group Variance | 0.15 | (0.39) |  | AIC |  |  |  | 2328.31 |  |
| Between-Group Variance |  |  |  | BIC |  |  |  | 2483.07 |  |
| Random Intercept (Participant) | 0.01 | (0.12) | 0.29 | R<sup>2</sup> (Total) |  |  |  | 0.94 |  |
|  |  |  |  | R<sup>2</sup> (Fixed Effects) |  |  |  | 0.94 |  |
| N Groups |  |  |  | ICC |  |  |  | 0.08 |  |
| Participant | 38 |  |  | RMSE |  |  |  | 0.38 |  |
| N Observations | 2280 |  |  |  |  |  |  |  |  |

**Table A5.** Coefficients describing the effect of *r* on *Z* by acoustic feature, averaged over experimental condition.

| Feature | Coefficient | SE | 95% CI | | $t$ | df | $p$ |
| --- | --- | --- | --- | --- | --- | --- | --- |
|  |  |  | Lower | Upper |  |  |  |
| Variation | 47.19 | 0.89 | 45.44 | 48.94 | 52.92 | 2253 | < 0.001 |
| Env Deriv | 33.80 | 0.53 | 32.75 | 34.85 | 63.19 | 2253 | < 0.001 |
| Centroid | 22.78 | 0.56 | 21.68 | 23.89 | 40.45 | 2253 | < 0.001 |
| Flux | 20.25 | 0.41 | 19.44 | 21.06 | 49.06 | 2253 | < 0.001 |
| Crest | 19.54 | 0.44 | 18.68 | 20.39 | 44.68 | 2253 | < 0.001 |
| Skew | 19.44 | 0.45 | 18.56 | 20.32 | 43.34 | 2253 | < 0.001 |
| Spread | 18.63 | 0.48 | 17.69 | 19.57 | 38.74 | 2253 | < 0.001 |
| Envelope | 16.97 | 0.36 | 16.27 | 17.67 | 47.41 | 2253 | < 0.001 |
| Slope | 15.46 | 0.38 | 14.72 | 16.20 | 40.79 | 2253 | < 0.001 |
| Decrease | 13.41 | 0.31 | 12.80 | 14.03 | 42.72 | 2253 | < 0.001 |

**Table A6.** Pairwise tests of estimated marginal means comparing the effect of *r* on *Z* by acoustic feature, averaged over experimental condition. *p*-values are Bonferroni-adjusted for multiple comparisons.

|  |  | Estimate | SE | 95% CI |  | *t* | DF | *p* | *d* | 95% CI |  |
| --- | --- | --- | --- | --- | --- | --- | --- | --- | --- | --- | --- |
|  |  |  |  | Lower | Upper |  |  |  |  | Lower | Upper |
| Env Deriv | Envelope | 16.83 | 0.63 | 15.59 | 18.06 | 26.74 | 2253 | 0.00 | 0.56 | 0.52 | 0.61 |
| Centroid | Envelope | 5.81 | 0.66 | 4.52 | 7.10 | 8.82 | 2253 | 0.00 | 0.19 | 0.14 | 0.23 |
| Spread | Envelope | 1.66 | 0.59 | 0.50 | 2.81 | 2.81 | 2253 | 0.22 | 0.06 | 0.02 | 0.10 |
| Skew | Envelope | 2.47 | 0.56 | 1.36 | 3.57 | 4.39 | 2253 | 0.00 | 0.09 | 0.05 | 0.13 |
| Crest | Envelope | 2.56 | 0.55 | 1.48 | 3.65 | 4.64 | 2253 | 0.00 | 0.10 | 0.06 | 0.14 |
| Slope | Envelope | -1.51 | 0.51 | -2.51 | -0.51 | -2.97 | 2253 | 0.14 | -0.06 | -0.10 | -0.02 |
| Decrease | Envelope | -3.56 | 0.46 | -4.47 | -2.65 | -7.67 | 2253 | 0.00 | -0.16 | -0.20 | -0.12 |
| Variation | Envelope | 30.21 | 0.95 | 28.36 | 32.07 | 31.94 | 2253 | 0.00 | 0.67 | 0.63 | 0.72 |
| Flux | Envelope | 3.28 | 0.53 | 2.23 | 4.32 | 6.16 | 2253 | 0.00 | 0.13 | 0.09 | 0.17 |
| Centroid | Env Deriv | -11.02 | 0.77 | -12.52 | -9.51 | -14.36 | 2253 | 0.00 | -0.30 | -0.34 | -0.26 |
| Spread | Env Deriv | -15.17 | 0.71 | -16.56 | -13.78 | -21.41 | 2253 | 0.00 | -0.45 | -0.49 | -0.41 |
| Skew | Env Deriv | -14.36 | 0.69 | -15.71 | -13.01 | -20.92 | 2253 | 0.00 | -0.44 | -0.48 | -0.40 |
| Crest | Env Deriv | -14.26 | 0.68 | -15.59 | -12.93 | -21.01 | 2253 | 0.00 | -0.44 | -0.49 | -0.40 |
| Slope | Env Deriv | -18.34 | 0.64 | -19.60 | -17.08 | -28.48 | 2253 | 0.00 | -0.60 | -0.64 | -0.56 |
| Decrease | Env Deriv | -20.39 | 0.61 | -21.58 | -19.19 | -33.49 | 2253 | 0.00 | -0.71 | -0.75 | -0.66 |
| Variation | Env Deriv | 13.39 | 1.01 | 11.40 | 15.37 | 13.21 | 2253 | 0.00 | 0.28 | 0.24 | 0.32 |
| Flux | Env Deriv | -13.55 | 0.66 | -14.84 | -12.26 | -20.53 | 2253 | 0.00 | -0.43 | -0.48 | -0.39 |
| Spread | Centroid | -4.15 | 0.73 | -5.59 | -2.72 | -5.68 | 2253 | 0.00 | -0.12 | -0.16 | -0.08 |
| Skew | Centroid | -3.34 | 0.71 | -4.74 | -1.95 | -4.70 | 2253 | 0.00 | -0.10 | -0.14 | -0.06 |
| Crest | Centroid | -3.25 | 0.70 | -4.62 | -1.87 | -4.62 | 2253 | 0.00 | -0.10 | -0.14 | -0.06 |
| Slope | Centroid | -7.32 | 0.67 | -8.64 | -6.01 | -10.95 | 2253 | 0.00 | -0.23 | -0.27 | -0.19 |
| Decrease | Centroid | -9.37 | 0.64 | -10.62 | -8.12 | -14.72 | 2253 | 0.00 | -0.31 | -0.35 | -0.27 |
| Variation | Centroid | 24.40 | 1.04 | 22.36 | 26.45 | 23.44 | 2253 | 0.00 | 0.49 | 0.45 | 0.54 |
| Flux | Centroid | -2.53 | 0.69 | -3.89 | -1.18 | -3.68 | 2253 | 0.01 | -0.08 | -0.12 | -0.04 |
| Skew | Spread | 0.81 | 0.65 | -0.46 | 2.08 | 1.25 | 2253 | 1.00 | 0.03 | -0.01 | 0.07 |
| Crest | Spread | 0.91 | 0.64 | -0.35 | 2.16 | 1.42 | 2253 | 1.00 | 0.03 | -0.01 | 0.07 |
| Slope | Spread | -3.17 | 0.60 | -4.35 | -1.99 | -5.27 | 2253 | 0.00 | -0.11 | -0.15 | -0.07 |
| Decrease | Spread | -5.22 | 0.56 | -6.32 | -4.11 | -9.24 | 2253 | 0.00 | -0.19 | -0.24 | -0.15 |
| Variation | Spread | 28.56 | 1.00 | 26.60 | 30.52 | 28.58 | 2253 | 0.00 | 0.60 | 0.56 | 0.65 |
| Flux | Spread | 1.62 | 0.62 | 0.40 | 2.84 | 2.60 | 2253 | 0.42 | 0.05 | 0.01 | 0.10 |
| Crest | Skew | 0.10 | 0.61 | -1.10 | 1.30 | 0.16 | 2253 | 1.00 | 0.00 | -0.04 | 0.04 |
| Slope | Skew | -3.98 | 0.58 | -5.11 | -2.85 | -6.92 | 2253 | 0.00 | -0.15 | -0.19 | -0.10 |
| Decrease | Skew | -6.03 | 0.54 | -7.08 | -4.97 | -11.23 | 2253 | 0.00 | -0.24 | -0.28 | -0.19 |
| Variation | Skew | 27.75 | 0.98 | 25.82 | 29.67 | 28.23 | 2253 | 0.00 | 0.59 | 0.55 | 0.64 |
| Flux | Skew | 0.81 | 0.60 | -0.36 | 1.98 | 1.35 | 2253 | 1.00 | 0.03 | -0.01 | 0.07 |
| Slope | Crest | -4.08 | 0.56 | -5.19 | -2.97 | -7.22 | 2253 | 0.00 | -0.15 | -0.19 | -0.11 |
| Decrease | Crest | -6.12 | 0.53 | -7.15 | -5.09 | -11.64 | 2253 | 0.00 | -0.25 | -0.29 | -0.20 |
| Variation | Crest | 27.65 | 0.98 | 25.73 | 29.57 | 28.30 | 2253 | 0.00 | 0.60 | 0.55 | 0.64 |
| Flux | Crest | 0.71 | 0.59 | -0.44 | 1.87 | 1.21 | 2253 | 1.00 | 0.03 | -0.02 | 0.07 |
| Decrease | Slope | -2.05 | 0.48 | -2.99 | -1.10 | -4.26 | 2253 | 0.00 | -0.09 | -0.13 | -0.05 |
| Variation | Slope | 31.73 | 0.95 | 29.86 | 33.60 | 33.25 | 2253 | 0.00 | 0.70 | 0.65 | 0.75 |
| Flux | Slope | 4.79 | 0.55 | 3.71 | 5.87 | 8.72 | 2253 | 0.00 | 0.18 | 0.14 | 0.23 |
| Variation | Decrease | 33.77 | 0.93 | 31.94 | 35.60 | 36.21 | 2253 | 0.00 | 0.76 | 0.72 | 0.81 |
| Flux | Decrease | 6.84 | 0.51 | 5.84 | 7.83 | 13.50 | 2253 | 0.00 | 0.28 | 0.24 | 0.33 |
| Flux | Variation | -26.94 | 0.97 | -28.83 | -25.04 | -27.85 | 2253 | 0.00 | -0.59 | -0.63 | -0.54 |

**Table A7.** Pairwise tests comparing the effect of *r* on *Z* by experimental condition, within acoustic feature. *p*-values are Bonferroni-adjusted for multiple comparisons.

| Feature | Contrast | Estimate | SE | 95% CI | | $t$ | df | $p$ | $d$ | 95% CI | |
| --- | --- | --- | --- | --- | --- | --- | --- | --- | --- | --- | --- |
|  |  |  |  | Lower | Upper |  |  |  |  | Lower | Upper |
| Envelope | Dutch Vocoded - English Vocoded | 4.07 | 1.23 | 1.65 | 6.50 | 3.31 | 214 | 0.017 | 0.45 | 0.18 | 0.72 |
| Env Deriv | Dutch Vocoded + Blurring - English Vocoded + Blurring | -8.73 | 2.25 | -13.16 | -4.29 | -3.88 | 214 | 0.002 | -0.53 | -0.80 | -0.26 |
| Spread | Dutch Vocoded + Blurring - English Vocoded + Blurring | -4.23 | 1.22 | -6.64 | -1.83 | -3.47 | 214 | 0.009 | -0.47 | -0.75 | -0.20 |

### Appendix A.6. Sensitivity to Speech Understanding and Spectral Degradation

For each feature, we first built a baseline model containing only a random intercept for participant, to which the addition of a fixed effect of spectral degradation or language was compared. If both main effects improve model fit (likelihood ratio test *p* = 0.05), we fit a model with both terms and/or their interaction against each single main effect.

**Figure A2.**
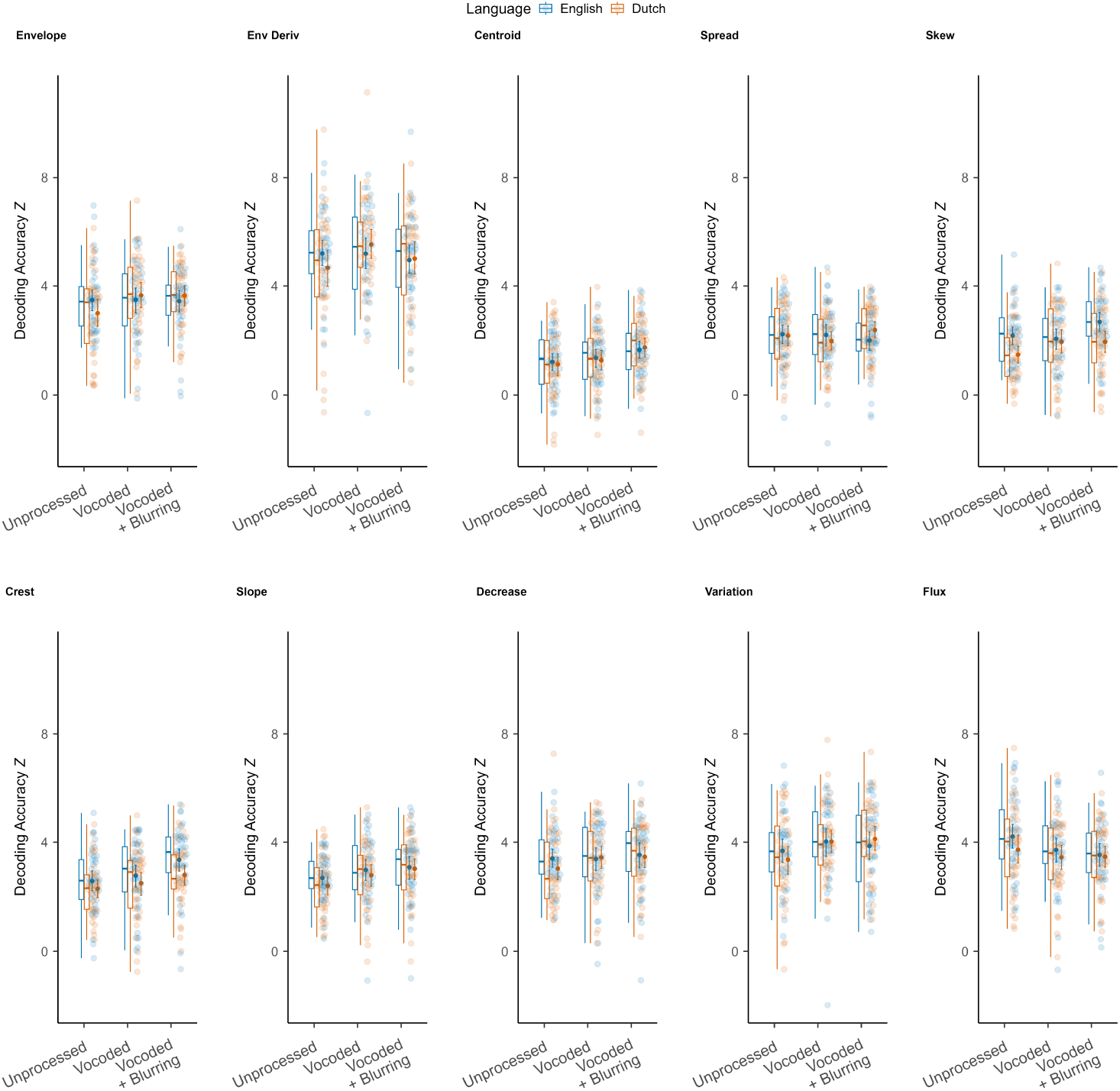
Acoustic features plotted by experimental condition. Box plots indicate the median and inter-quartile range of the distribution. Individual data points are pooled over participant and experimental condition. Solid markers represent the mean and error bars show the 95% confidence intervals of the mean.

**Table A8.**
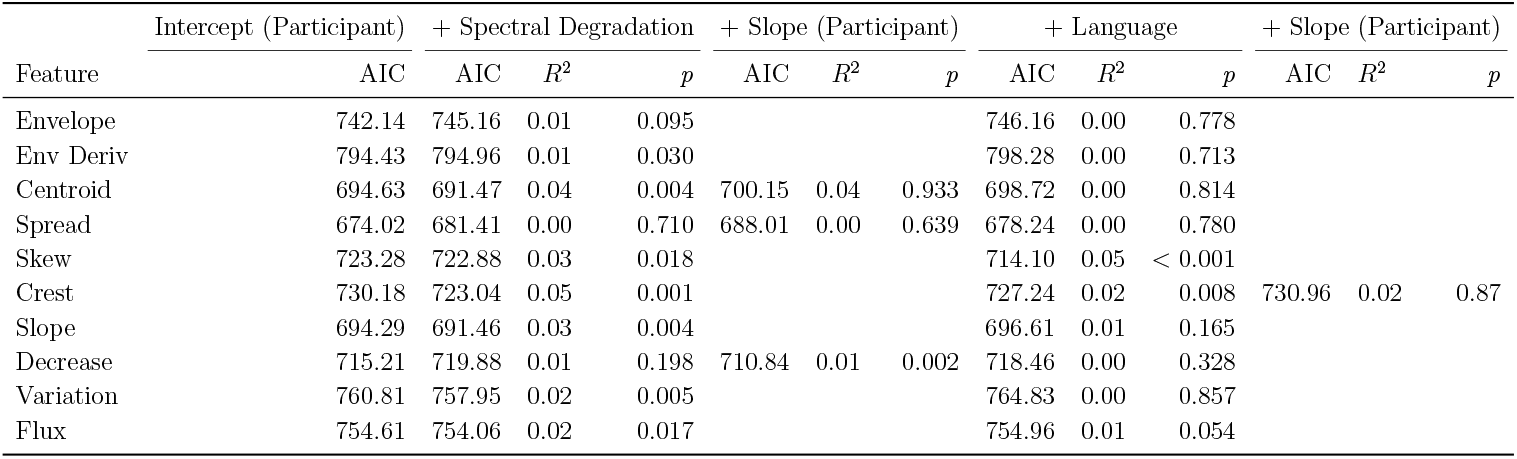
Comparison of linear mixed effect models of *Z* by acoustic feature testing the fixed effects of Spectral Degradation and Language. Each fixed effect is compared to the baseline model containing only an intercept for Participant. Missing values are where the model did not converge due to an unsupported random effects structure.

**Table A9.**
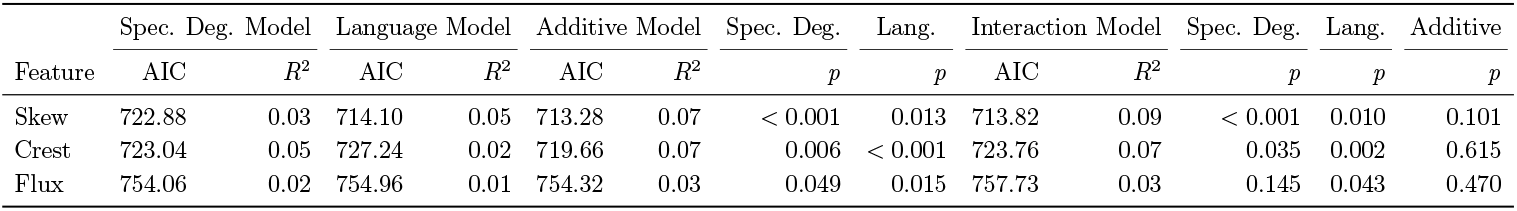
Comparison of linear mixed effect models of *Z* by acoustic feature testing the addition and interaction between Spectral Degradation and Language. The full interaction model is compared to models containing only main effects.

**Table A10.** Model formula: *Z ∼* (1 | Participant). Standardized parameters were obtained by fitting the model on a standardized version of the dataset. 95% Confidence Intervals (CIs) and p-values were computed using a Wald t-distribution approximation.

| Parameter | Estimate | 95% CI | | $t$ | df | $p$ | Std Est | 95% CI | |
| --- | --- | --- | --- | --- | --- | --- | --- | --- | --- |
|  |  | Lower | Upper |  |  |  |  | Lower | Upper |
| (Intercept) | 3.45 | 3.13 | 3.78 | 21.34 | 37.00 | 0.00 | 0.00 | -0.24 | 0.24 |
| Random Effects | Variance | SD | ICC | Model Metrics |  |  |  |  |  |
| Within-Group Variance | 1.11 | (1.06) |  | AIC |  |  |  | 742.14 |  |
| Between-Group Variance |  |  |  | BIC |  |  |  | 752.43 |  |
| Random Intercept (Participant) | 0.81 | (0.9) | 0.65 | R<sup>2</sup> (Conditional) |  |  |  | 0.42 |  |
|  |  |  |  | R<sup>2</sup> (Marginal) |  |  |  | 0.00 |  |
| N Groups |  |  |  | ICC |  |  |  | 0.42 |  |
| Participant | 38 |  |  | RMSE |  |  |  | 0.98 |  |
| N Observations | 228 |  |  |  |  |  |  |  |  |

#### Appendix A.6.1. Spectral Envelope Model

**Table A11.**
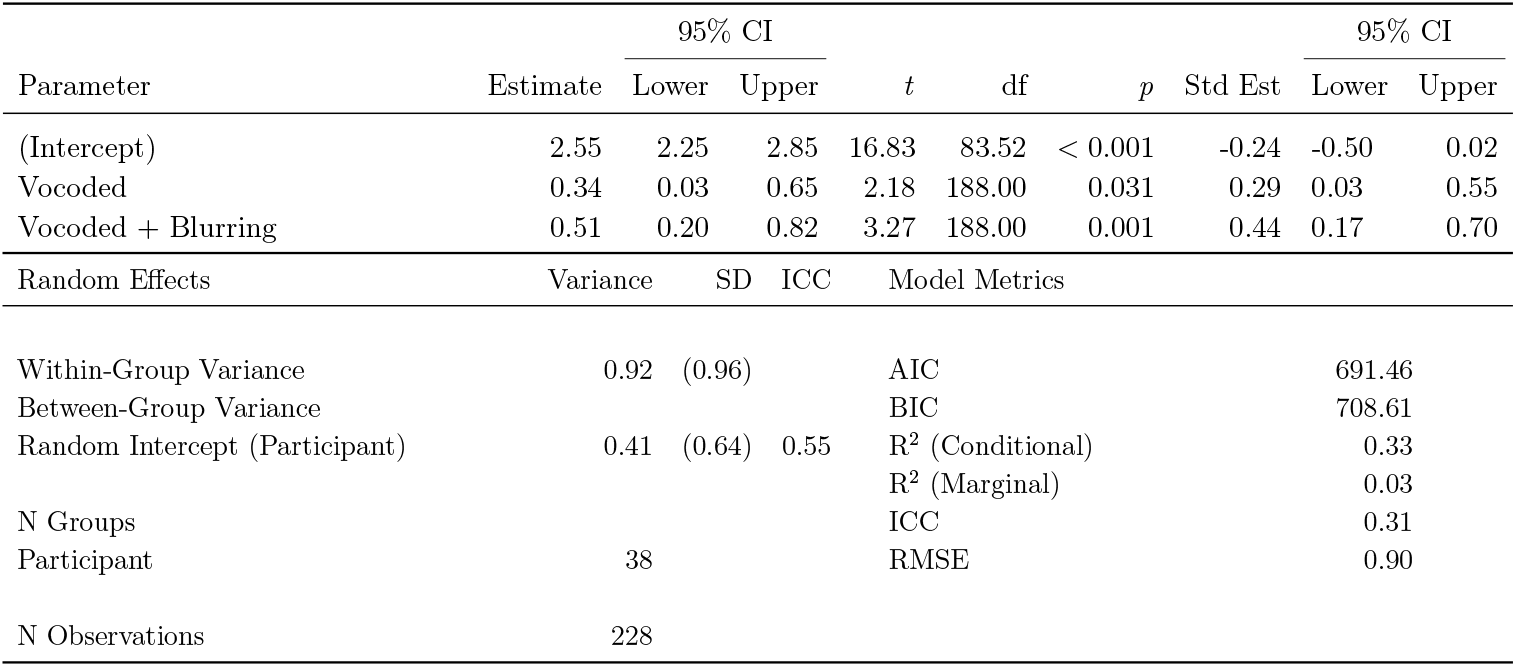
Model formula: *Z ∼* Spectral Degradation + (1 | Participant). Standardized parameters were obtained by fitting the model on a standardized version of the dataset. 95% Confidence Intervals (CIs) and p-values were computed using a Wald t-distribution approximation.

| Parameter | Estimate | 95% CI |  | *t* | df | *p* | Std Est | 95% CI |  |
| --- | --- | --- | --- | --- | --- | --- | --- | --- | --- |
|  |  | Lower | Upper |  |  |  |  | Lower | Upper |
| (Intercept) | 2.55 | 2.25 | 2.85 | 16.83 | 83.52 | < 0.001 | -0.24 | -0.50 | 0.02 |
| Vocoded | 0.34 | 0.03 | 0.65 | 2.18 | 188.00 | 0.031 | 0.29 | 0.03 | 0.55 |
| Vocoded + Blurring | 0.51 | 0.20 | 0.82 | 3.27 | 188.00 | 0.001 | 0.44 | 0.17 | 0.70 |
| Random Effects | Variance | SD | ICC | Model Metrics |  |  |  |  |  |
| Within-Group Variance | 0.92 | (0.96) |  | AIC |  |  |  | 691.46 |  |
| Between-Group Variance |  |  |  | BIC |  |  |  | 708.61 |  |
| Random Intercept (Participant) | 0.41 | (0.64) | 0.55 | R<sup>2</sup> (Conditional) |  |  |  | 0.33 |  |
|  |  |  |  | R<sup>2</sup> (Marginal) |  |  |  | 0.03 |  |
| N Groups |  |  |  | ICC |  |  |  | 0.31 |  |
| Participant | 38 |  |  | RMSE |  |  |  | 0.90 |  |
| N Observations | 228 |  |  |  |  |  |  |  |  |

#### Appendix A.6.2. Spectral Slope Model

**Table A12.**
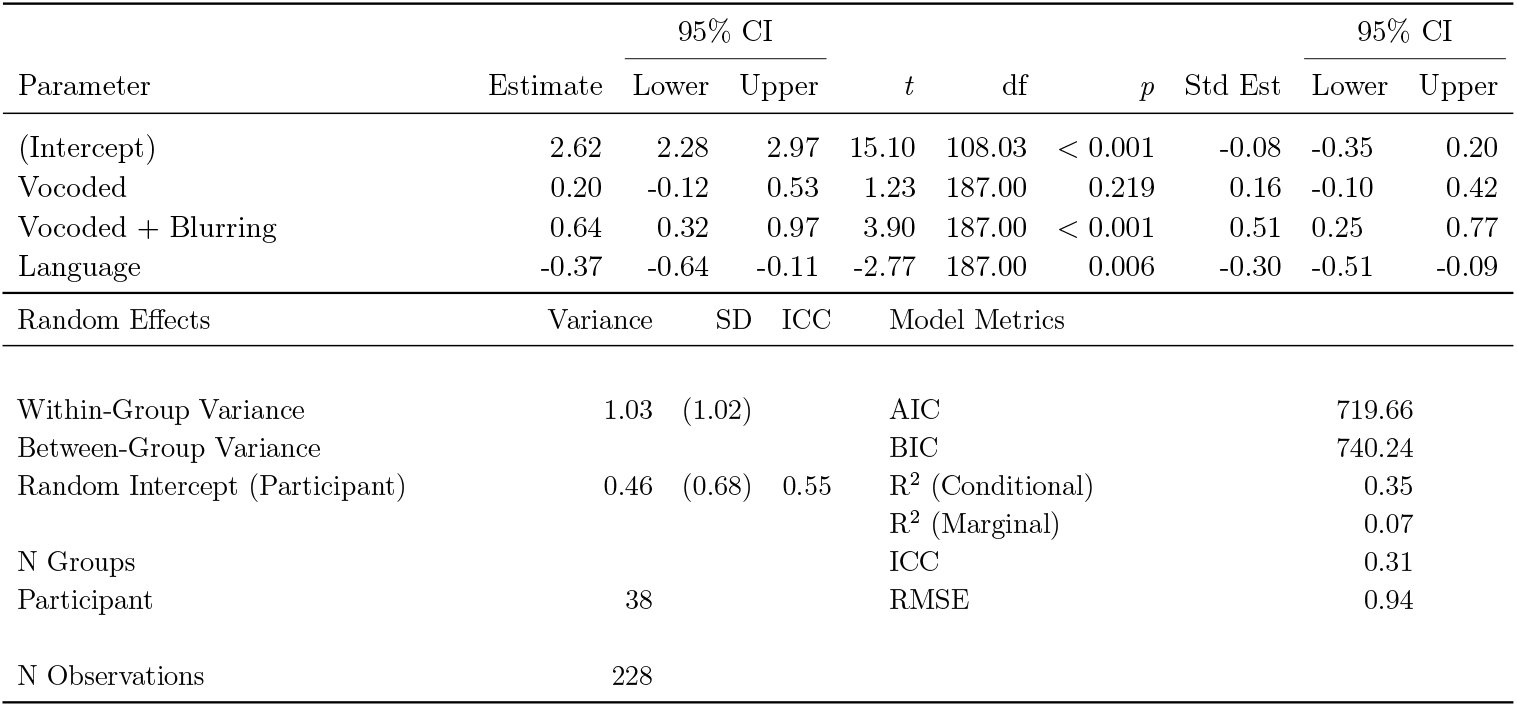
Model formula: *Z ∼* Spectral Degradation + Language + (1 | Participant). Standardized parameters were obtained by fitting the model on a standardized version of the dataset. 95% Confidence Intervals (CIs) and p-values were computed using a Wald t-distribution approximation.

| Parameter | Estimate | 95% CI | | $t$ | df | $p$ | Std Est | 95% CI | |
| --- | --- | --- | --- | --- | --- | --- | --- | --- | --- |
|  |  | Lower | Upper |  |  |  |  | Lower | Upper |
| (Intercept) | 2.62 | 2.28 | 2.97 | 15.10 | 108.03 | < 0.001 | -0.08 | -0.35 | 0.20 |
| Vocoded | 0.20 | -0.12 | 0.53 | 1.23 | 187.00 | 0.219 | 0.16 | -0.10 | 0.42 |
| Vocoded + Blurring | 0.64 | 0.32 | 0.97 | 3.90 | 187.00 | < 0.001 | 0.51 | 0.25 | 0.77 |
| Language | -0.37 | -0.64 | -0.11 | -2.77 | 187.00 | 0.006 | -0.30 | -0.51 | -0.09 |
| Random Effects | Variance | SD | ICC | Model Metrics |  |  |  |  |  |
| Within-Group Variance | 1.03 | (1.02) |  | AIC |  |  |  | 719.66 |  |
| Between-Group Variance |  |  |  | BIC |  |  |  | 740.24 |  |
| Random Intercept (Participant) | 0.46 | (0.68) | 0.55 | R<sup>2</sup> (Conditional) |  |  |  | 0.35 |  |
|  |  |  |  | R<sup>2</sup> (Marginal) |  |  |  | 0.07 |  |
| N Groups |  |  |  | ICC |  |  |  | 0.31 |  |
| Participant | 38 |  |  | RMSE |  |  |  | 0.94 |  |
| N Observations | 228 |  |  |  |  |  |  |  |  |

#### Appendix A.6.3. Spectral Crest Model

**Table A13.**
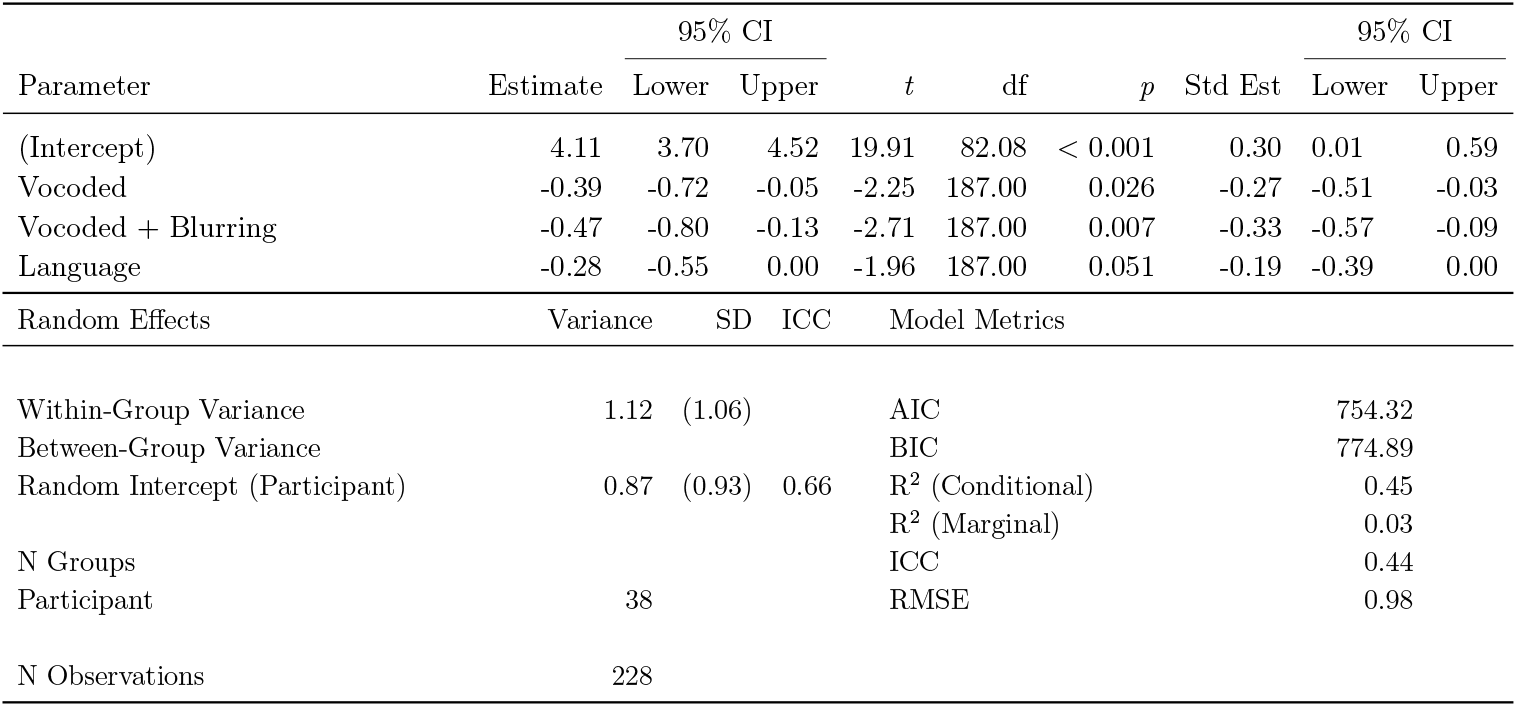
Model formula: *Z ∼* Spectral Degradation + Language + (1 | Participant). Standardized parameters were obtained by fitting the model on a standardized version of the dataset. 95% Confidence Intervals (CIs) and p-values were computed using a Wald t-distribution approximation.

#### Appendix A.6.4. Spectral Flux Model

*‡* Note that identifying the acoustic correlates of phonetic information in speech is a topic of debate and ongoing research within linguistics, see [28, 29] for discussion.

